# Evolutionary contingency’s impact on laboratory evolution of *Escherichia coli* under fluctuating environments

**DOI:** 10.1101/598995

**Authors:** Ximo Pechuan, Daniel Biro, Maryl Lambros, Aviv Bergman

## Abstract

1

The adaptation of biological organisms to fluctuating environments is one major determinant of their structural and dynamical complexity. Organisms have evolved devoted adaptations to ensure the robust performance of physiological functions under environmental fluctuations. To further our understanding of particular adaptation strategies to different environmental fluctuations, we perform laboratory evolution experiments of *Escherichia coli* under three temperature fluctuation regimes alternating between 15°C and 43°C. Two of these regimes are determined by the population’s growth, while the third regime switches stochastically. To address evolutionary contingencies, the experiments are performed on two lineages departing from different genetic backgrounds. The two lineages display distinct evolutionary trajectories, demonstrating dependency on the starting strain’s genetic background. Several genes exhibit a high degree of parallelism, suggesting their potential adaptive nature. The growth increase of the representative clones from each final population relative to their ancestor at 15°C and 43°C demonstrated no correlation between both temperatures, insinuating an absence of a strong trade-off between these two temperatures. Some had a growth rate decrease at 15°C unless exposed to a 43°C epoch, indicating some degree of internalization of the structure of the environment fluctuations. The phenotypic response of the evolved clones at 15°C and 43°C was assessed by a phenotype array method. The resulting responses reveal a general tendency to move closer to the phenotypic response of our starting strains at the optimum of 37°C. This observation expands the documented restorative responses, even when facing complex environmental conditions.

**Author Summary:** Laboratory evolution experiments have been widely employed to test hypotheses from evolutionary theory. To assess the dynamics of adaptation under environmental fluctuations, we evolved 24 *Escherichia coli* populations under different regimes of temperature switching between 15°C and 43°C for about 600 generations. At the final point of the evolution experiment, the evolved populations were genome sequenced and clones were isolated and sequenced for phenotypic characterization. Fitness measurements revealed adaptation to both environmental conditions and some strains internalized the environmental fluctuation. Array phenotypic measurements showed that the majority of evolved strains tended to restore the phenotypic signature of the perturbed environments to that of the optimal temperature condition. This observation expands the documented restorative responses, even when facing complex environmental conditions.

## 3 Introduction

Fluctuating environmental conditions are conjectured to play a major role in evolution (Frank 2011, Levins 1968). The complex dynamical nature of both biotic and abiotic factors are thought to be one of the evolutionary drivers of the high ecological diversity of species that have populated our biosphere over time (Fuentes & Ferrada 2017). The environmental complexity, besides the con-natural potential of life (Ruiz-Mirazo et al. 2008), may explain the seemingly open ended nature of the evolutionary process. More concretely, biological populations respond to environmental conditions by a variety of adaptive strategies that require devoted molecular mechanisms. For example, environmental complexity promotes the evolution of robustness mechanisms, phenotypic plasticity or life-cycle strategies (Kaneko 2012, Pigliucci 2001).

The particularities, scale and predictability of environmental fluctuations play a fundamental role in the specific ways biological organisms adapt to different selective pressures that vary dynamically or spatially (Botero et al. 2015). Upon encountering environmental diversity, some hypothesize that organisms face the choice of adapting to one environment, namely becoming specialists, or to adapt to a subset of the possible environments, namely becoming generalists. The ultimate reason for these two limiting strategies has been conjectured to be the existence trade-offs between the adaptive responses to different environmental conditions. The granularity and the predictability of environmental changes has also been hypothesized to determine the evolution of a particular adaptive strategy (Brown & Pavlovic 1992, Futuyma & Moreno 1988). Thus, it is suggested that predictable temporal environmental variation can lead to the evolution of phenotypic plasticity and internalization of this environmental cycle. Less predictable environmental variation can lead to stochastic phenotypic switching, bet-hedging, or specialization to the most frequent environment (Beaumont et al. 2009, Thattai & van Oudenaarden 2004).

Although both ecological and paleontological data can be used to assess these hypotheses, adaptive laboratory evolution experiments provide a controlled setting where tenets about the evolutionary process can be tested on model organisms whose biology is known in great detail (Elena & Lenski 2003, Herron & Doebeli 2013). Even though the results derived from these experiments might be hard to extend to the whole biosphere, they provide us with a rich phenomenology that can potentially be understood at the molecular level and give new insights that can be used to construct better theoretical frameworks. One of the most commonly used laboratory evolution organisms is *Escherichia coli* and many questions regarding the dynamics of adaptation to different environmental stresses have been formulated with the set-up that Lenski and co-workers established as the pioneers of modern era experimental evolution (Barrick et al. 2009, Lenski et al. 1991). This set-up has been extensively used together with genome sequencing to explore a manifold of adaptive scenarios involving different degrees of complex environmental conditions to which the ancestral strain was not adapted to (Satterwhite & Cooper 2015).

Using *Escherichia coli* as the organism and temperature as the environmental variable is the ideal set-up to investigate the effects of time-scale, determinism, and amplitude of environmental fluctuations on the evolutionary dynamics. Temperature has been widely chosen as a target environmental variable in studies of experimental evolution of micro-organisms because of its global implications on an organism’s physiological state and growth, and its easy manipulation (Riehle et al. 2001, Rudolph et al. 2010, Sandberg et al. 2014, Tenaillon et al. 2012). Consequently, there is extensive knowledge of mutations that are to be expected and their degree of parallelism within and across experimental evolution studies for certain temperatures. Most *Escherichia coli* adaptive laboratory evolution studies focus on temperatures closer to the upper thermal boundary, usually around 42°C. There are far fewer studies that focus on the adaptation at colder temperatures than the optimal temperature of 37°C, and none of them go far below ambient temperature. Few experimental evolution studies controlling temperature variations have also been carried out (Bennett et al. 1992, Ketola et al. 2013, Kishimoto et al. 2010, Saarinen et al. 2018), however they have not extensively evaluated the consequences of different environmental fluctuation scenarios as is addressed in this work.

In this work, we make use of laboratory evolution of *Escherichia coli* in fluctuating temperatures to address whether extreme environmental fluctuations lead to the development of adaptive trade-offs (understood as specialist strategies) under different fluctuation regimes. While *Escherichia coli* is well adapted to grow at 37°C and has shown its potential for adapting to a variety of environmental challenges, we wanted to assess its ability to adapt to two alternating environmental conditions close to the temperature niche boundary (15°C and 43°C). Given the different physiological state required to address extreme low and high temperatures, we conjectured that a trade-off might limit the adaptation to both conditions. As the scale and predictability of the environmental change would also have an effect in the resulting adaptive strategies, we tested two environments that fluctuated in a deterministic fashion with two different generation time scales and an additional randomly fluctuating condition. This choice to dynamically switch environmental treatments is predicated on the different adaptive responses that are conjectured to dominate in these different environments. The two treatments determined by the optical density of the culture, in principle, should favor the evolution of a generalist strategy as the variants that adapt to both environments will be favored. These two treatments differ in the optical density that triggers the temperature change and thus in the number of generations that population experiences at a given temperature segment allowing us to assess the effects of the generation timescales in the types of adaptive responses evolved. Additionally, given their predictable nature can be potentially internalized by the organism. The stochastic treatment serves as a control where the number of generations that experience a certain environment does not depend on the population density. In this scenario, given that the 43°C has a higher growth rate than the 15°C, a potential strategy can be to adapt primarily to the 43°C environment and see 15°C as a perturbation, thus favouring the appearance of specialists. To investigate the dependence of the evolutionary outcome with the genetic background of the starting point, the starting populations were derived from REL606 and REL607 strains adapted to the media at their optimal temperature of 37°C, and mutations previously associated with adaptation to glucose minimal media were observed. Thus, the starting points of the two lineages allowed us to address the effects of their mutational background on their adaptation to each fluctuating temperature condition by comparing the two lineages’ adaptive trajectories.

## 4 Results

### 4.1 Mutational Background of the Starting Points

The initial adaptation period to the media of the original REL606 and REL607 strains led to a series of mutations in our starting REL606 and REL607 populations in addition to the original *araA* (D92G) mutation (see Table S1). Our starting REL607 population also had a mutation in the exonuclease V gene *recD* (V10A) that has been found to be neutral with respect to Lenski’s original REL606 strain and already present in the Lenski’s long term experimental evolution (LTEE) strains (Phillips & Wilson 2016). Our starting REL606 population acquired two mutations with potential impact on the fitness of the strain: a large deletion in the *rbs* operon involved in ribose metabolism and uptake genes, and a mutation in the translation initiation factor gene (*infB)*. Deletions in the *rbs* operon have been previously reported to evolve in glucose media laboratory evolution experiments with a high degree of parallelism (Phillips & Wilson 2016, Tenaillon et al. 2016) due to the combination of being selectively advantageous and having a high mutation rate mediated by the nearby IS transposable element (Cooper et al. 2001). The *infB* mutation has also been observed in similar experiments (Barrick et al. 2009) and could also be related to adaptation to the media. Accompanying these fixed mutations were a couple of deletions, one per line, that were almost fixed in our starting populations. Our starting REL606 population carried a large deletion starting in *cybB*. The deletion starting at *cybB* was mediated by an insertion element and affected a total of eight genes, including the the *cybB* gene coding for a cytochrome, components of a toxin antitoxin system (*hokB, mokB)*, transcription regulators, components of chemotactic systems, and glycan synthesis proteins. Deletions surrounding *hokB/mokB* mediated by IS elements have been reported to occur in Lenski’s LTEE strains and in mutation accumulation strains indicating the presence of a mutational hotspot (Tenaillon et al. 2016, Woods et al. 2006). For our REL607 starting population, a deletion of Δ122 bp in the hypothetical protein *ECB 02621* was also almost fixed. In both of our starting populations, mutations in the gene *nadR* were also observed which are common in experimental evolution of these strains with glucose as a carbon source (Phillips & Wilson 2016). Two other mutations were observed at a low frequency; an intergenic mutation in the *yhjY/tag* in REL607 and a nonsynonymous mutation in the *uxaB* in REL606. In our laboratory evolution experiments, our starting points were two heterogeneous populations resulting from a pre-adaptation process. To ensure that the observed mutations in the evolved strains were indeed not present as low frequency mutants in the starting populations, we sequenced the starting populations again at a high coverage (to an average of 6000). Additional very low frequency mutants were detected in this sequencing run but none of them were carried over to evolved populations. The starting populations’ distinct mutational backgrounds enabled us to assess the contingency of the evolutionary trajectory on the initial conditions.

### 4.2 Mutational Analysis of the Final Populations

The genes affected by mutations present in our heterogeneous populations at the final time points for each environmental regime and strain are summarized in Figure 1. A more detailed set of tables describing the specific characteristics of the mutations can be found in the supplementary information (see Table S3). From the mutational pattern in our final populations, we observe that nonsynonymous mutations and small indels are the most frequent types of mutations. Large amplifications and deletions mediated by transposable elements are also common and could have potentially dramatic adaptive consequences. Overall there seems to not be a population accumulating substantially more mutations than the average, meaning that mutations appear and rise in our populations at about the same rate for all three environmental treatments (slow, fast, and random).

**Figure 1:**
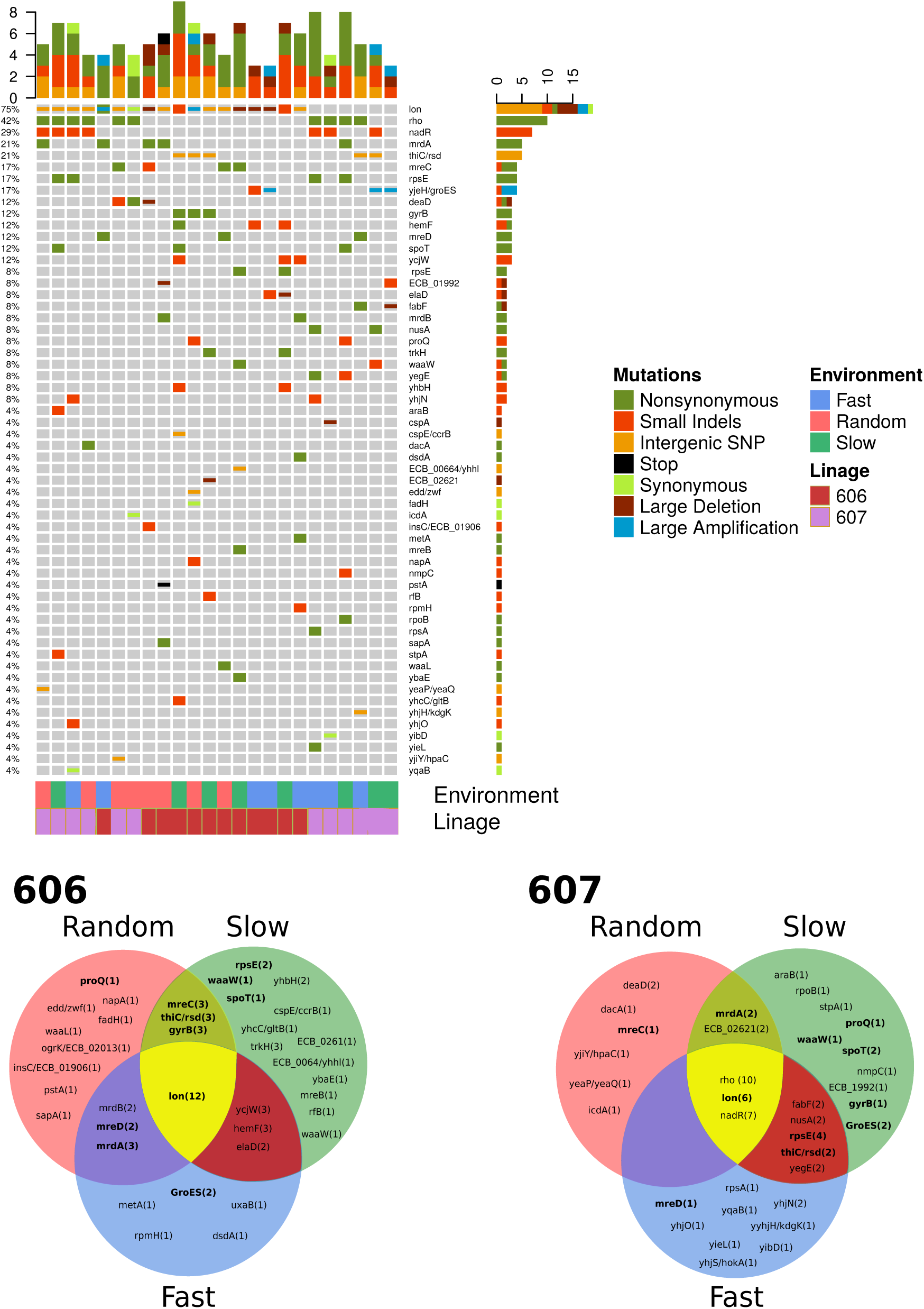
Mutational composition of our final populations. The upper panel shows the mutational signature of each population. Mutations are coarse grained at the gene level. Large deletions and amplifications are counted once and collapsed to the repeatable gene. The lower panel shows the Venn Diagram of this information separated by strain. The numbers in parentheses indicate the number of our populations with a mutation in the gene. The genes in bold had mutants in both our REL606 and REL607 final populations. Note, we consider a large indel when it is greater than 100 bp.

Of all the mutations observed, two were highly repeatable at the level of the genomic region affected and were present in almost all of our populations irrespective of the environmental treatments; mutations in the ATP-dependent DNA-RNA helicase *rho* and mutations in the intergenic region of the *lon* heat shock protease (see Figure 1 Upper Panel). The high degree of repeatability of these mutations in most of our final populations indicated that they may be the main contributors to the growth rate increase observed in our adaptive evolution experiment. It is important to note that the parallelism was also present at the molecular level of the particular mutation in some instances of the intergenic region of *lon* and to a great extent in *rho*. Mutations in *rho*, like the ones that evolved in our experiment have also been consistently appearing with a high degree of repeatability in adaptive evolution experiments of *Escherichia coli* to high temperatures (Tenaillon et al. 2012), suggesting a crucial role of the termination factor in this environmental condition. The REL606 final populations <u>lack</u> mutations in the *rho* gene indicating that the adaptive benefit of these mutations is strongly contingent on the genetic background, which could have conditioned this distinct evolutionary trajectory seen in our experiments.

Most of the remaining mutations are present in more than one population, but do not have the high degree of parallelism observed in the two formerly described mutations. Some of these remaining mutations can arguably be associated with a particular environmental treatment. The two deterministic treatments, slow and fast, can be linked to mutations in the regulatory genes: *rpsE, nusA, ycjW, yhbH*. The ribosomal protein gene *rpsE* is also mutated with high parallelism, affecting the same amino acid, and appears related to the deterministic regimes in both strains. The presence of this mutation in the deterministic environments suggest that it might be important for cold adaptation. The *nusA* non-synonymous substitutions are also related to *rho* termination as the *nusA* gene encodes a transcription termination/antitermination factor. It is not surprising that only our REL607 final populations have mutations in *nusA* as only these REL607 final populations have mutations in *rho* further supporting the potential dependency on the genomic background.

Mutations genes associated with cell envelope, such as *mrd* and *mre* genes, appear predominantly in the random environmental regime. Most of these cell envelope related genes have also been observed to mutate in laboratory evolution experiments using glucose media (Tenaillon et al. 2016) this could indicate further adaptation to the media rather than to our temperature regimes. Similarly, there are mutations in several metabolic genes, such as thiamine synthesis gene *thiC* and *spoT*, which are related to adaptation to glucose media rather than the temperature conditions (Phillips & Wilson 2016, Tenaillon et al. 2016).

Additionally, other mutations are present at high frequencies in particular populations. A salient set of these mutations affect heat shock proteins, like the chaperonin gene *groEL*. Two ways of potentially increasing chaperonin proteins expression arose, one through large genomic region amplifications and the other through a specific deletion in the regulatory region of the *groEL* gene. The large genomic amplifications affect a large number of proteins involved in the heat shock response, potentially having a great adaptive value for our populations. Similar to the small deletion in *groEL*, a intergenic region mutation in this chaperonin gene has been previously reported in high temperature adaptation (Yama et al. 2015).

Overall, a wide variety of mutations emerged in relatively few generations with high degree of parallelism even to the molecular detail. More interestingly, the genomic amplification of the heat shock genes was also convergent. It is important to note that a most of the big deletions and amplifications were mediated by mobile elements which evidences the big role they play in increasing the adaptability of the strains that carry them.

### 4.3 Clone characterization

#### 4.3.1 Mutations

For each of our final populations, we isolated and sequenced a set of clones for further characterization of their phenotypes. The representative clones were consistent with their parental populations and their dominant mutations (see Tables 1-3). The clonal isolate from the F607-1 population had a discrepancy with its parental population in terms of the mutational composition. The lower sequencing coverage of this particular population could explain this discrepancy in terms of the clonal isolate belonging to a low frequency mutant missed by the sequencing. The sequenced clones allowed us to support the presence of large amplifications and deletions suggested by the heterogeneous population data. The clonal sequencing data also helped to further discern the synteny relations between mutations observed in the heterogeneous populations with similar frequencies.

**Table 1:** Mutations found in the clones isolated from our final populations evolved in the random environmental treatment.

| Genes | Function | Mutation Annotation | Strains |
| --- | --- | --- | --- |
| **Ancestral:** |  |  |  |
| rbsD-[rbsR] | Ribose Metabolism | IS150-mediated $\Delta$ 5,874 bp at 3,894,997 | R606-1,R606-2,R606-3,R606-4 |
| cybB-[ydcG] | Multiple Genes | IS1-mediated Δ6,496 bp at 1,460,928 | R606-1,R606-2,R606-4 |
| infB | Translation | V85D (GTC→GAC) | R606-1,R606-2,R606-3,R606-4 |
| nadR | NAD metabolism | Coding (658/1233 nt) +G at 4,616,186 | R606-1,R606-2,R606-3,R606-4 |
| araA | Arabinose metabolism | D92G (GAC→GGC) | R607-1,R607-2,R607-3,R607-4 |
| recD | Exonuclease | V10A (GTT→GCT) | R607-1,R607-2,R607-3,R607-4 |
| yhjY/tag | Biofilm formation/DNA repair | Intergenic (-42/-116) G→A at 3,643,434 | R607-1,R607-2,R607-4 |
| nadR | NAD metabolism | E367* stop (GAG→TAG) | R607-3 |
| ECB_02621 | Unannotated | Coding $\Delta$ 122 bp at 2,792,849 | R607-3 |
| **Regulatory:** |  |  |  |
| rho | Termination factor | I15N (ATC→AAC) | R607-2,R607-3,R607-4 |
| rho | Termination factor | H295Y (CAT→TAT) | R607-1 |
| polA/yihA | DNA Synthesis/Cell Division | Intergenic (+245/+135) Δ2 bp at 4,028,294 | R606-1 |
| **Cold Shock:** |  |  |  |
| deaD | Ribosome Assembly | Coding (1223/1890 nt) +66bp at 3,241,872 | R607-3 |
| **Heat Shock:** |  |  |  |
| insL2/lon | Transposase/heat shock protease | Intergenic (-48/-108) C → T at 430,835 | R607-3,R607-4 |
| insL2/lon | Transposase/Heat shock protease | Intergenic (-92/-64) G → T at 430,879 | R606-2,R606-4, R607-1 |
| insL2/lon | transposase/heat shock protease | Intergenic IS mediated Δ1,350 bp at 429,504 +24 bp at 430,879 | R606-1 |
| insL2-[ppiD] | Multiple genes | Amplification at [430,618-434,631] | R606-3 |
| **Metabolism:** |  |  |  |
| nadR | NAD metabolism | Coding (165/1233 nt) Δ1bp at 4,615,693 | R607-1,R607-2,R607-4 |
| yjiY/hpaC | Membrane Protein/Reductase | Intergenic (-347/+46) T/A at 456,960 | R607-3 |
| edd/zwf | Glycolysis | Intergenic (-30/+205) A→C at 1,913,210 | R606-3 |
| thiC/rsd | Thiamine Biosynthesis/ Stationary Phase | Intergenic (-166/+68) A→T at 4,175,876 | R606-3 |
| pstA | Phosphate Uptake | W264* (TGG→TGA) | R606-2 |
| napA | Nitrate Reductase | Coding (1510/2487 nt) Δ1bp at 2,251,232 | R606-3 |
| **Cell Envelope:** |  |  |  |
| dacA | Cell Wall Synthesis | K76T (AAA→ACA) | R607-1 |
| mrdA | Cell Wall Synthesis | Y397S (TAC→TCC) | R607-4 |
| mrdA | Cell Wall Synthesis | V175I (GTC→ATC) | R606-2 |
| mreC | Cell Shape Regulation | Coding (1077/1104 nt) +C at 3,326,793 | R606-1 |
| mreD | Cell Shape Regulation | P28L (CCG→CTG) | R606-4 |
| **Others:** |  |  |  |
| ogrK-[ECB 02013] | Multiple genes | Coding $\Delta$ 22,146 bp at 2,100,308 | R606-2 |
| proQ | RNA Chaperone | Coding (431435/699) nt $\Delta$ 5 bp | R607-2 |
| proQ | RNA Chaperone | Coding (333-345/699 nt) $\Delta$ 13bp at 1,893,767 | R606-3 |
| gyrB | DNA Coiling | R732H (CGC→CAC) | R606-3 |

**Table 2:** Mutations found in the clones isolated from our final populations evolved in a fast, deterministic environmental treatment.

| Gene | Function | Mutation Annotation | Strains |
| --- | --- | --- | --- |
| **Ancestral:** |  |  |  |
| rbsD-[rbsR] | Ribose Metabolism | IS150-mediated $\Delta$ 5,874 bp at 3,894,997 | F606-1, F606-2, F606-3, F606-4 |
| cybB-[ydcG] | Multiple Genes | IS1-mediated Δ6,496 bp at 1,460,928 | F606-1, F606-2, F606-3, F606-4 |
| infB | Translation | V85D (GTC→GAC) | F606-1, F606-2, F606-3, F606-4 |
| nadR | NAD metabolisim | Coding (658/1233 nt) +G at 4,616,186 | F606-1, F606-2, F606-3, F606-4 |
| araA | Arabinose metabolism | D92G (GAC→GGC) | F607-1,F607-2,F607-3,F607-4 |
| recD | Exonuclease | V10A (GTT→GCT) | F607-1,F607-2,F607-3,F607-4 |
| yhjY/tag | Biofilm formation/DNA repair | Intergenic (-42/-116) G→A at 3,643,434 | F607-2 |
| nadR | NAD metabolism | E367* stop (GAG→TAG) | F607-1,F607-3 |
| ECB_02621 | Unannotated | Coding $\Delta$ 122 bp at 2,792,849 | F607-1,F607-3 |
| **Regulatory:** |  |  |  |
| rho | Termination factor | I15N (ATC→AAC) | F607-2,F607-4 |
| rho | Termination factor | I15S (ATC→AGC) | F607-3 |
| rpsE | Ribosomal Protein | G91S (GGT→AGT) | F607-3 |
| rpsE | Ribosomal Protein | C112S (TGC→AGC) | F607-2 |
| ycjW | Transcriptional Regulator | Coding (177-179/999 nt) IS150 (-) +3 bp | F606-1 |
| yhnH | Ribosome Regulator | Coding (273/288 nt) +6bp at 3,281,680 | F606-1 |
| **Heat Shock:** |  |  |  |
| insL2/lon | transposase/heat shock protease | Intergenic (-24/-132) G→A at 430,811 | F606-1 |
| insL2/lon | transposase/heat shock protease | Non-coding and intergenic (-68/-88) Δ1350 bp at 429505 | F606-2 |
| yjeH/groES | transporter/chaperonin | Intergenic (-200/-74) Δ3 bp at 4,349,333 | F606-2 |
| [insL-2]-[ppiD] | Multiple genes | Amplification at [430689-435601] | F606-3 |
| [dsbD]-efp/ecnA | Multiple genes | Amplification at [4343517-4354989] | F606-4 |
| yjdN-[insG] | Multiple genes | Amplification at [4304004-4485407] | F607-1 |
| **Metabolism:** |  |  |  |
| amn/yeaN | AMP degradation/Chaperone Related | Intergenic (+260/-83) T→C at 1,994,774 | F607-1 |
| yieL | Xylanase | W29R (TGG→CGG) | F607-3 |
| hemF | Oxidase | Coding (657-683/900 nt) Δ27bp at 2,481,400 | F606-2 |
| kup/insJ-5 | Potassium transporter/IS150 protein | Intergenic (+6/-50) +G at ,893,551 | F606-1 |
| ygcQ | Flavoprotein | T161T (ACC→ACA) | F606-3 |
| nadR | NAD metabolism | Coding (165/1233 nt) Δ1bp at 4,615,693 | F607-2,F607-3 |
| **Cell Envelope:** |  |  |  |
| yfgA | Cell Wall Shape | Coding (841-877/1014 nt) $\Delta$ 37 bp at 2,562,787 | F607-1 |
| yhjN | Cellulose Synthase Regulator | Coding (2237-2245/2340 nt) Δ9bp at 3,619,855 | F607-2,F607-3 |
| mrdA | Cell Wall Synthesis | N428D (AAC→GAC) | F606-3 |
| **Others:** |  |  |  |
| ECB_01992 | Unannotated | Amplification coding (185/216 nt) | F606-2 |
| elaD | Deubiquitinating protease | Coding (734/1212 nt) $\Delta$ 1 bp at 2,329,952 | F606-4 |
| [yhjS]-hokA | Multiple genes | IS 150 meadiated $\Delta$ 23682 bp at 3,627,409 | F607-4 |

**Table 3:** Mutations found in the clones isolated from our final populations evolved in a slow, deterministic environmental treatment.

| Gene | Function | Mutation Annotation | Strains |
| --- | --- | --- | --- |
| **Ancestral:** |  |  |  |
| rbsD-[rbsR] | Ribose Metabolism | IS150-mediated $\Delta$ 5,874 bp at 3,894,997 | S606-1, S606-2, S606-3, S606-4 |
| cybB-[ydcG] | Multiple Genes | IS1-mediated Δ6,496 bp at 1,460,928 | S606-1, S606-2, S606-3, S606-4 |
| infB | Translation | V85D (GTC→GAC) | S606-1, S606-2, S606-3, S606-4 |
| nadR | NAD metabolim | Coding (658/1233 nt) +G at 4,616,186 | S606-1, S606-2, S606-3, S606-4 |
| araA | Arabinose metabolism | D92G (GAC→GGC) | S607-1,S607-2,S607-3,S607-4 |
| recD | Exonuclease | V10A (GTT→GCT) | S607-1,S607-2,S607-3,S607-4 |
| yhjY/tag | Biofilm formation/DNA repair | Intergenic (-42/-116) G→A at 3,643,434 | S607-1,S607-3,S607-4 |
| nadR | NAD metabolism | E367* stop (GAG→TAG) | S607-2 |
| ECB_02621 | Unannotated | Coding $\Delta$ 122 bp at 2,792,849 | S607-2 |
| **Regulatory:** |  |  |  |
| rho | Termination factor | I15N (ATC→AAC) | S607-3,S607-4 |
| nusA | Elongation factor | T72A (ACC→GCC) | S607-1 |
| rpsE | Ribosomal Protein | G91C (GGT→TGT) | S607-4 |
| rpsE | Ribosomal Protein | G91A (GGT→GCT) | S606-4 |
| **Heat Shock:** |  |  |  |
| cadC/pheU-[yjeM] | Multiple Genes | Amplification at [4,341,178-4,362,612] | S607-1 |
| [fxsA]-[yjeP] | Multiple Genes | Amplification at [4,347,401-4,366,500] | S607-2 |
| insL2/lon | transposase/heat shock protease | Intergenic (-24/-32) G→A at 430,811 | S607-3 |
| insL2/lon | transposase/heat shock protease | Intergenic (-48/-108) C → T at 430,835 | S606-1 |
| insL2/lon | transposase/heat shock protease | Intergenic Δ1350bp at 429,505 | S606-4 |
| insL2/lon | transposase/heat shock protease | Intergenic (-69/-87) Δ1bp at 430,856 | S606-3 |
| **Cold Shock:** |  |  |  |
| fabF-pabC | Cold Fatty Acid Synthesis | Coding $\Delta$ 650 bp at 1,167,222 | S607-2 |
| **Metabolism:** |  |  |  |
| ribC | riboflavin metabolism | F157L (TTT→TTG) | S607-2 |
| hemF | Oxidase | Coding (621/900 nt) $\Delta$ 1 bp at 2,481,364 | S607-2 |
| hemF | Oxidase | W124R (TGG→AGG) | S606-2 |
| prpE/codB | propionyl-CoA synthase/cytosine transporter | Intergenic (+198/-37) $\Delta$ 2bp at 327,440 | S607-3 |
| potA/pepT | ABC transporter/Peptidase | Intergenic (-132/-118) A→G at 1,201,096 | S606-2 |
| trkH | Potassium transporter | T20P (ACC→CCC) | S606-2 |
| trkH | Potassium transporter | S349A (TCA→GCA) | S606-1 |
| thiL | Thiamine Biosynthesis | L34F (CTC→TTC) | S606-3 |
| nadR | NAD metabolism | Coding (165/1233 nt) Δ1bp at 4,615,693 | S607-1,S607-3 |
| **Cell Envelope:** |  |  |  |
| nmpC | Membrane Porin | Pseudogene (421/606 nt) T→G at 547,858 | S607-4 |
| impF/asns | Membrane Porin/tRNA synthetase | Intergenic (-91/+411) 1 $\Delta$ bp at 1,004,265 | S607-3 |
| mreB | Cell Wall Synthesis | G117S (GGC→AGC) | S606-4 |
| mreC | Cell Wall Synthesis | Coding (49/1104 nt) (CACCGCCAG)1→2 at 3,327,821 | S606-2 |
| **Other:** |  |  |  |
| ECB_02621 | Unannotated | Coding (645-705/849 nt) Δ61bp at 2,793,153 | S606-1 |
| elaC-yfbM | Multiple Genes | Coding Δ3382bp at 2,328,714 | S606-3 |
| elaD | Deubiquitinating protease | Coding (734/1212 nt)) Δ1bp at 2,329,959 | S607-1 |
| yegE | Motility | Coding (1456-1462/3306 nt) $\Delta$ 7 bp at 2,077,927 | S607-4 |

#### 4.3.2 Temperature adaptation

In order to evaluate the adaptation of our populations to the experimental temperatures, we measured the growth rate at 15°C and 43°C for each representative clone relative to their ancestral starting strains. We also measured these growth rates for the original Lenski strains relative to each other and our starting strains to detect changes due to the mutational background acquired during the adaptation to the media period.

The original Lenski strains showed no significant fitness differences under any of the temperatures studied further supporting the claim that the *Ara* mutation is neutral under a wide range of environmental conditions (see Table S9). After the initial media adaptation period, our two starting strains gained a significant fitness difference at 37°C relative to the original Lenski strains they were derived from. Moreover, our starting REL606 strain had a significantly higher growth rate at 37°C than our starting REL607 strain. The metabolic nature of the background mutations accumulated by our starting REL606 strain suggests adaptation to the media as a plausible explanation for the growth difference (see Table S2). Both starting strains also showed a growth rate increase at 43°C suggesting that adaptation to the media at the optimal temperature can also lead to changes in the response to other temperatures which justifies the adaptation to the media period. The REL 606 starting strain also showed an increase in growth rate at 15°C. The background mutations of this strain confers an unexpected advantage at the cold temperature probably derived from a better utilization of glucose as the most salient mutations in this strain involve the deletion of many of the ribose operon genes. Nevertheless, the descendants of our starting REL606 strain further adapted to the cold temperature, demonstrating that the background mutations did not exhaust the adaptive potential to this environmental condition in our experiment.

For the representative clones of our final populations (see Table 4), the general observed pattern is the absence of a trade-off between the adaptation to 15°C and 43°C. Both temperatures show a fitness increase in most of our clonal strains, and there is no significant correlation between the fitness increase of the two experimental temperatures (Pearson correlation, *r* = −0.16, p-value = 0.457). In both our populations, increases in fitness of the lineages evolved under random environmental fluctuations mostly occur at the 43°C temperature. A possible explanation could be the nature of the environmental treatment itself. The independence of the bacterial growth from the temperature switching time is a plausible explanation for this adaptive response in the random treatment. The random environmental fluctuation posed an adaptive problem where the 43°C condition could be regarded as the adaptive target and 15°C as a perturbation. The reason for this asymmetry is in the intrinsically higher growth rate at the 43°C environment. Most strains evolved in the deterministic environmental treatments, which the temperature switch depends on population growth, showed fitness increases in both the hot and cold temperatures. Surprisingly, two of our clonal strains experiencing a deterministic treatment showed a significant fitness decrease at 15°C (see Table 4). It is important to note that this fitness decrease only occurred after an acclimation pass at 15°C. If the strain experienced the hot temperature in the acclimation pass, then the fitness was neutral with respect to its starting strain (see Supplementary Figure 2). Some of the mutations carried by these strains, like the deletion of *fabF*, could have deleterious effects in 15°C but, according to our observations, be neutral in the context of a previous exposure to the 43°C environment. This is indicative that the deterministic environment has been internalized to some degree by these populations.

**Table 4:** Relative fitness of our clonal strains measured by comparison of their growth rate to their respective starting strain’s growth rate. Colored in green the significant fitness increases and in red the significant fitness decreases in the measurement setting.

| Strain | T=15°C | T=43°C |
| --- | --- | --- |
| Random 607-1 | 1.09* | 1.76* |
| Random 607-2 | 0.94 | 1.34* |
| Random 607-3 | 1.09 | 1.19* |
| Random 607-4 | 0.99 | 1.33* |
| Fast 607-1 | 1.26* | 1.34* |
| Fast 607-2 | 1.21* | 1.45* |
| Fast 607-3 | 1.27* | 1.12* |
| Fast 607-4 | 1.22* | 1.09* |
| Slow 607-1 | 1.77* | 1.33* |
| Slow 607-2 | 0.25* | 1.37* |
| Slow 607-3 | 1.33* | 1.66* |
| Slow 607-4 | 1.27* | 1.57* |
| Random 606-1 | 0.94 | 1.03 |
| Random 606-2 | 1.09 | 1.20* |
| Random 606-3 | 1.00 | 1.14* |
| Random 606-4 | 1.02 | 1.19* |
| Fast 606-1 | 0.98 | 1.24* |
| Fast 606-2 | 0.86* | 1.44* |
| Fast 606-3 | 1.03 | 1.36* |
| Fast 606-4 | 1.40* | 1.18* |
| Slow 606-1 | 1.52* | 1.35* |
| Slow 606-2 | 0.66* | 1.42* |
| Slow 606-3 | 1.18* | 1.24* |
| Slow 606-4 | 1.12* | 1.18* |

\*Two sample t-test, heterocedastic, P-val < 0.05

**Figure 2:**
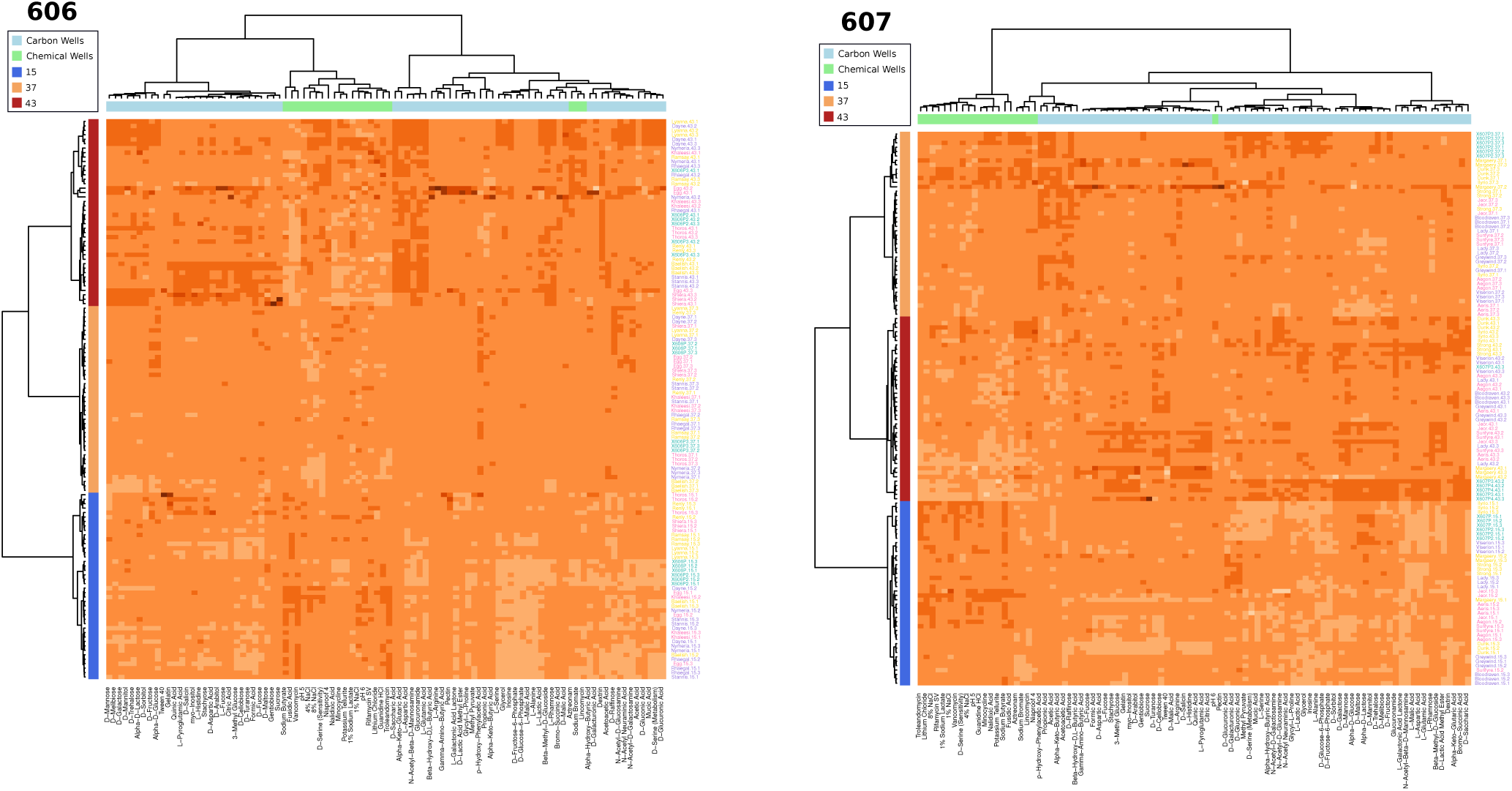
Hierarchical clustering of the Biolog phenotypic data. The rows are colored by the experimental temperature. The columns are colored by the type of well of the Biolog plate. Finally, the rows are accompanied by the strain name colored by the treatment (random as light pink, fast as purple, and slow as yellow).

#### 4.3.3 Evolved Phenotypic Response

In order to understand the consequences of the acquired mutations on the tendencies shown by the phenotypic response of the strains relative to their ancestors, we performed a set of phenotypic assays at the three relevant temperatures of 15°C, 37°C and 43°C. The phenotypes evaluated corresponded to metabolic and chemical sensitivity assays that our strains were not exposed to during our laboratory evolution experiments. This phenotypic evaluation allows us to study the consequences of the evolutionary adaptation to our conditions in other regions of the ecological niche. Additionally, the resulting phenotypic signature can also be used as a proxy to study the hypothesized restorative effects of the accumulated mutations during adaptive laboratory evolution experiments, which is conjectured to bring the phenotypic signature closer to that of the unstressed environment (Hug & Gaut 2015). Here, we extend this hypothesis to the evolutionary response in complex environmental treatments.

Clustering of the data resulting from the phenotypic assays showed that the primary discriminating factor in the phenotypic response of the different strains was the temperature treatment (see Figure 2). For both lineages, the cluster trees show two primary splits corresponding to the 15°C data and, the 43°C and 37°C data. The latter then splits into the two respective leafs for these two temperatures. This implies that the 37°C environment is more similar to the 43°C environment than to the 15°C, and that the 15°C condition is also possibly more stressful as it appears to be further from the optimum in terms of the phenotypic response. It is important to note that some clustering of the phenotypes was observed for our REL607 clones, separating the chemical sensitivity assays from the metabolic tests. Clustering by temperature fluctuation regime is apparent, but not significant enough to be conclusive. In general, the replicates tend to cluster together justifying taking averages in the posterior analyses.

We performed dimensionality reduction using principal component analysis (PCA) on the measurements resulting from the phenotypic assays, in order to eliminate the effects of potential correlation between the phenotypes measured by the Biolog plates (which are not necessarily designed to obtain orthogonal phenotypes). Because of the different mutational background of our starting strains, we kept our REL606 and REL607 clones from our final populations separated in the analysis. Looking at the PCA of our starting strains, we see that both strains can be distinguished further justifying the separate analysis (see Figure S3). Four principal components are informative of the patterns in the starting strains data. The first principal component discriminates between the different temperatures. The second principal component discriminates between the optimal temperature and the other two temperatures. Finally, the fourth principal component discriminates between the strains. There also seems to be interaction between the strains and the temperature as the local structure of the strain factor is different for each temperature cluster.

The same pattern can be seen in the PCA of the evolved lineages with the first principal component distinguishing the temperatures and the second principal component distinguishing the optimal from the perturbed temperatures (see Figure 3). When assessing the direction of the evolved phenotype response by using the classification and methodology described in (Hug & Gaut 2015), the restorative effect is more prominent in the 607 lineage in the first two principal components (see Table S13). The 606 lineage shows little restoration for the 43° environment while more of the phenotypic response has tended to the optimal for the 15° environment. Even though some of the clones exhibited reinforcement, both lineages showed substantial amount of restoration to its starting strain’s phenotypic response for the first two retained principal components. The fourth and third principal component showed more reinforced and even novel responses, nevertheless these account for a smaller fraction of the variance. In general, our data demonstrates that there is a general tendency towards the restoration of the phenotypic signature exhibited in the optimal environment even in the face of a more complex adaptive challenge.

**Figure 3:**
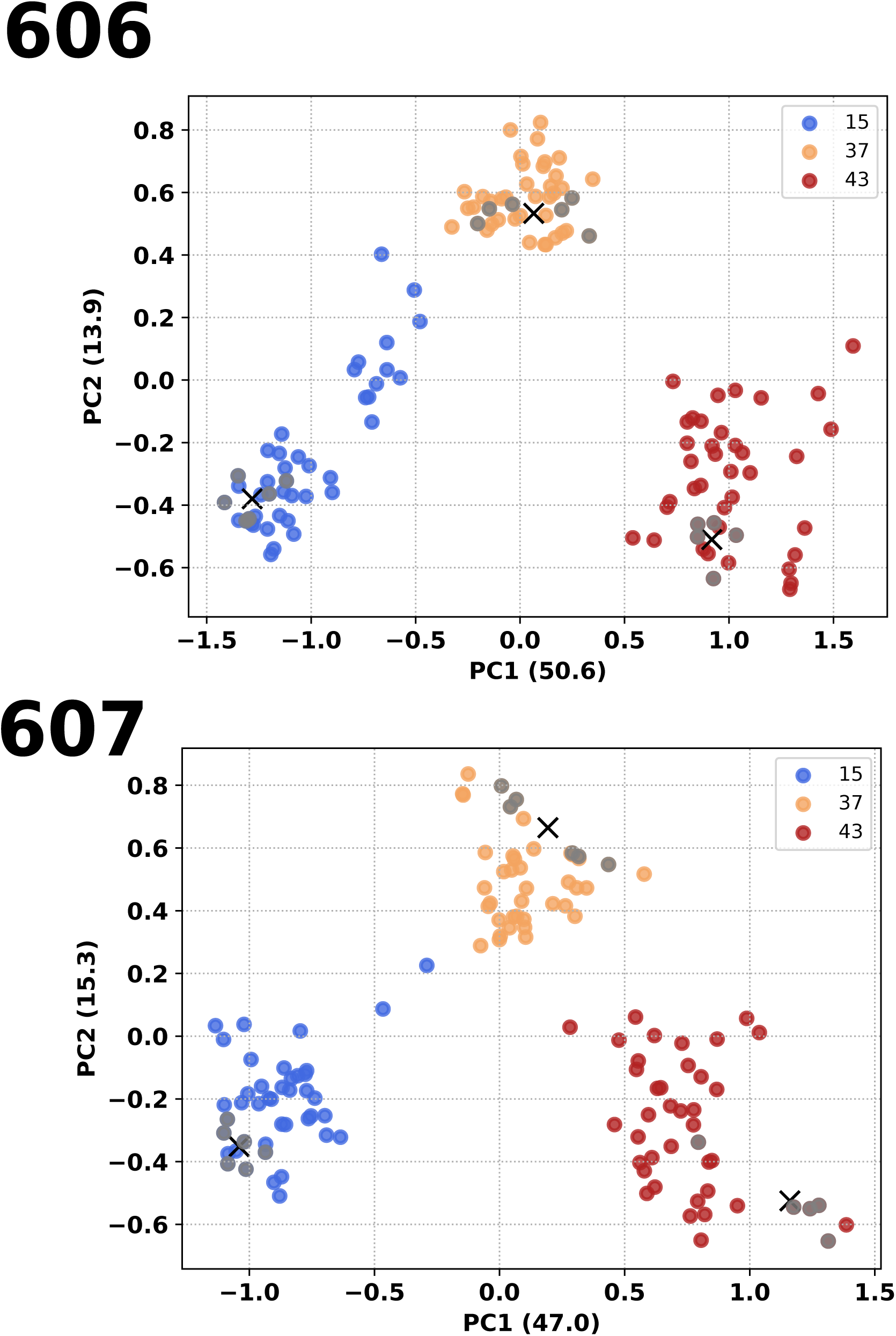
Principal component analysis of the phenotypic fingerprint at the three temperatures of our clones of our final populations, together with their respective starting strain. The colors indicate the temperature treatments of the Biolog plates (15°C, 37°C, and 43°C) and our reference starting strain’s response at those temperatures. The crosses mark the centroids of the clusters of our starting strain at each of the three temperatures. The axes shows the amount of variance explained by each of the components.

## 5 Discussion

The implications of our laboratory evolution experiment in the current understanding of the adaptation of *Escherichia coli* to fluctuating temperatures can be discussed in terms of the mutational observations, the fitness gains in both environments related to the temperature fluctuation regime, and the adaptation impacts on the phenotypic signatures to such conditions in relation to the unstressed state. The mutation sets that evolved in our experiment are solid candidates for explaining the evolved clones’ increase in growth observed in both of the stressful temperatures because of their high degree of repeatability. Additional reconstruction experiments need to be carried out in order to unveil any potential epistatic interactions between the mutations. In addition, transcriptomic and proteomic analyses are needed to clarify the transcriptional and translational consequences of these mutations. Nevertheless, the potential consequences of the mutations can be discussed on the basis of our experimental observations and the current state of the literature.

In our experiment, the split between our two evolved lineages regarding the acquisition of a mutation in the termination factor *rho* can help clarify the molecular mechanisms that make *rho* mutants so pervasive in *Escherichia coli* adaptive laboratory evolution experiments at high temperatures. In this line of research, there is recent evidence that suggests the role of the mutations in *rho*, as observed in our experiment, is to restore the gene expression pattern of the unstressed condition (González-González et al. 2017). Yet, the distinct evolutionary trajectories observed for our two different lineages points towards an alternative explanation for the high parallelism of the *rho* mutations. Our results suggest that the background mutations of our starting strains conditioned the observed disjoint evolutionary trajectories. These include, the *rbs* deletion, *infB* non-synonymous mutations, and large *cybB* deletion. The *rbs* deletion and the other low frequency mutations observed in our starting REL606 strain affect metabolism or DNA repair, which makes them poor candidates for epistatically preventing the appearance of the *rho* mutations observed in our REL607 final populations and clones (see Table S1 and Figure 1 Upper Panel). However, there are two potential candidates in our starting REL606 strain that could prevent *rho* mutations from occurring in our REL606 final populations and clones.

The first potential candidate would be the mutation in the translation initiation factor *infB*. In the *Escherichia coli* genome this gene is in the operon of *nusA*, an important gene component of *rho*- dependent termination. Furthermore, *infB* has altered protein levels in *rho* inhibited cells (Cardinale et al. 2008). Thus, we hypothesize there may be a functional relationship between *infB* and *rho*, such that the *infB* background mutation in our starting REL606 strain may have similar functional consequences to our observed *rho* mutations.

The second possible explanation could be that the *cybB* deletion in our starting REL606 strain is sufficient to compensate for a the potential fitness benefit of the *rho* mutation in our REL607 final populations and clones. This deletion contains the toxin/antitoxin genes, *hokB* and *mokB*. We conjecture that the reason why *rho* mutations are so common in laboratory evolution experiments is related to the expression of genes with toxic effects on *Escherichia coli* genome, like *hokB* and *mokB*. Many toxic genes are terminated in a *rho*-dependent manner under normal conditions (Cardinale et al. 2008). Therefore, in a stressful environment the termination abilities of *rho* could be impacted such that some of these toxic genes are suddenly expressed, which would have a fitness cost. We hypothesize that the *cybB* deletion observed in our starting REL606 strain might have ablated some of these toxic elements potentially under the control of *rho*. In any event, the mutational background of the starting point in laboratory evolution experiments must be taken seriously when considering the adaptive value of mutations in any laboratory evolution experiment. All adaptive laboratory experiments are contingent on the evolutionary history of the starting point reflected on its genomic architecture.

Another salient feature of our experimental results is that a significant fraction of the mutations affect components of the proteostatic machinery involved in the heat-shock response (Nonaka 2006). Some of these mutations occurred through large genome amplifications that presumably could lead to an increase in the expression of these proteins. The major genes affected were those of the chaperonin GroEL/ES and the heat shock protease Lon. Lon is an ATP-dependent protease that is induced upon heat shock. The Lon protease participates in cellular proteostasis by degrading misfolded proteins preventing aggregate formation (Rosen et al. 2002). This protease is also involved in some regulatory interactions as some of its clients are regulatory elements (Aertsen & Michiels 2005, Dopazo et al. 1987, Van Melderen & Aertsen 2009). Affecting the expression levels of *lon* will presumably have an effect on protein degradation. The complex relationships of the protein quality control system and its fitness effects allow for the formulation of reasonable hypotheses for *lon* mutations observed in our experiment causing both an increase or decrease in expression (Bershtein et al. 2013, Cho et al. 2015). The temperature changes in our experiment may promote protein misfolding and the consequent accumulation of aggregates in the cytoplasm. An increased activity of *lon* could compensate for the permanent presence of these aggregates. On the other hand, a decrease in *lon* activity could also have an adaptive explanation in permanent stress conditions by allowing for the misfolded clients to be refolded by the other arm of the protein quality control system. It has been shown that defective *lon* mutants can rescue temperature sensitive *rpoD* mutants by increasing the amount of the sigma factor due to lower degradation in the high temperature conditions (Grossman et al. 1983). In order to adapt to constant high temperature stresses it may be preferred to lower the expression of the Lon protease in order to increase the amounts of other proteins whose activity has been reduced as consequence of the temperature increase.

Regarding the mutations observed in our final clonal strains that could confer fitness benefits at the cold temperature, the lack of studies in adaptation to this cold temperature condition make it difficult to assess the role that the observed mutations play in the fitness gains at this temperature (Dragosits & Mattanovich 2013)(see Table 4). Nevertheless, we hypothesize that the mutations in the regulatory elements *rpsE* and *ycjW* are potential candidates given their high repeatability in our REL607 and REL606 final clones (see Figure 1 Upper Panel). The many membrane and cell wall mutations observed could also play a role in adaptation to cold temperatures, but some of them have been observed in laboratory evolution experiments carried out at optimal temperature (Barrick et al. 2009). A way to address this issue would be to perform a laboratory evolution experiment at low temperatures in the spirit of the many examples of adaptation at high temperatures (Sandberg et al. 2014). This would also help discern a caveat of our study, which is the difficultly in establishing what adaptive limits do each of our two temperatures impose on each other. For instance in our experiment, although we observe many *rho* mutants that are common in laboratory evolution experiments, we have only one instance of the the also frequently observed *rpoB* mutations. The *rho* I15N mutation is reported to increase the high temperature niche boundary in adaptive laboratory evolution experiments (Rodríguez-Verdugo et al. 2014) with no discernible effect on the growth at lower temperatures. In contrast, the *rpoB* mutations have been demonstrated to exhibit fitness trade-offs at lower temperatures. The fact that *rpoB* mutations produce a niche shift rather than the niche expansion could potentially explain its absence in our experiment.

Another interesting feature of our experiment is the appearance of mutations that significantly lower the fitness in the cold environment, like the *fabF* and *deaD* deletions. However, our results suggest that these same mutations can be neutral in that environment as long as there is a previous epoch of the hot environment (see Table 4 and Figure S2). Our results highlight that the complexity of the environmental treatment needs to be taken into account when aiming to draw general conclusions about the trade-offs set by the mutational basis of a particular adaptation strategy.

Lastly, our experiment considered for first time the evaluation of the directionality of the phenotypic response of *Escherichia coli* strains simultaneously evolved in two different environmental conditions. We observe that, predominantly, our evolved strains’ responses to both temperatures move closer to the unperturbed state of its starting strain’s response to 37°C (see Figure 4). This restorative response is relevant in light of the potential limitations to adaptation that the dynamical varying nature of our experimental regimes pose. Measuring an array of phenotypes, generated by use of Biolog assays here, is a first step towards the characterization of the niche breadth and the adaptive effects of a particular environmental treatment on that breadth. Using laboratory evolution experiments smartly, we can also aim to better understand effects on niche breadth as driven by evolutionary contingencies.

## 6 Material and Methods

### 6.1 Experimental Evolution

The experiment was seeded with isolated clones from *Escherichia coli* REL606 and REL607 strains, which are derivatives of *Escherichia coli* B, provided by the Lenski laboratory. The strains were chosen for their wide use in experimental evolution studies. The REL607 strain harbors a neutral marker that renders it unable to metabolize arabinoise (*Ara*^*-*^) and distinguishable from the REL606 strain in tetrazolium and arabinose (TA) plates. Clones were isolated from a single colony of the respective strain and then propagated daily in M9 minimal media (Difco) supplemented with glucose 4 *g/L* for one month at 37°C in order to adapt them to the experimental media conditions. These cultures were transferred by taking a 100*μL* of culture and transferring to a new tube containing 20 *mL* of fresh media. From these two adapted starting cultures that harbored different mutational backgrounds from each other (see Table S2), 24 tubes of fresh media were seeded with either the REL606 or REL607 starting culture in replicates of four for each of the three temperature fluctuation treatments (2 strains× 3 treatments×4 replicates = 24) for a total of 12 seeded cultures for each of the two starting cultures. The three temperature fluctuation treatments were comprised of oscillations of the culture temperature between 15°C and 43°C (see Supplementary Figure 1). The time for a culture tube to reach the desired temperature after a temperature switch was approximately 30 minutes, however temperature acclimation to within two degrees of the target temperature was achieved in less than one generation. Of these three treatments, two were dependent on the optical density of the culture and one was random. The first optical density (OD) dependent treatment, termed “slow” or “S”, corresponded to a slow fluctuating environment with changes of temperature every time the strains reached an *OD*_830_ of 0.25, roughly half of their carrying capacity in our conditions. The second OD-dependent treatment, termed “fast” or “F”, corresponded to a faster fluctuating environment in generation times switching between the two temperatures at *OD*_830_ 0.15, 0.30, 0.45 respectively. The random treatment, termed “random” or “R”, sampled a random time for the culture to remain at a particular temperature from a uniform distribution between 3 to 5 hours at 43°C and 5 to 15 hours at 15°C. Using the *OD*_830_ measurements, growth phase was closely monitored, and bacteria were consistently transferred near the desired point in late exponential growth phase as the bacteria neared saturation at the end of each treatment cycle. Since each of the treatments contained four REL606 and four REL607 strains, it allowed us to follow the transfer schema in (Lenski et al. 1991) by alternating both strains providing means for testing for cross-contamination. Every transfer was performed by taking a 100*μL* of culture and transferring to a new tube containing 20 mL of fresh media. All cultures were incubated using a commercial batch culture BioSan LV Personal Bioreactor RTS1-C (Riga, Latvia), which controlled temperature to within 0.1°C. *OD*_830_ measurements were taken by the RTS-1C at 4 minute intervals. The instruments allowed us to program our deterministic fluctuations as a function of the optical density measurements and the random temperature fluctuation treatment. All tubes were cultured at 2000 rpm with 1 second reverse spinning intervals to ensure oxygenation. The experiment lasted around 6 months totalling an average of 600 generations.

### 6.2 Population Archive

Approximately every week, each evolving population was glycerol stocked. After transferring 100*μ*L of the culture after each treatment cycle to fresh media, the remaining culture was centrifuged for 10 minutes at 4000 rpm to form a pellet, and the excess liquid was removed. The pellet was then re-suspended in 1 mL of fresh M9 media and 1 mL of glycerol (Sigma) 20% to a final glycerol concentration of 10%. The samples were then placed directly in a freezer at −80 °C for long term storage. Whenever the bacteria were glycerol stocked they were also cultured in tetrazolium and arabinose (TA) M9 agar plates at 37°C overnight checking for the correct colored phenotype to detect potential external and cross contamination, using the protocols of (Lenski et al. 1991).

### 6.3 Generations Elapsed during the Adaptive Evolution Experiments

The number of generations for each of the 24 adapted strains was calculated by transforming the optical density measurements, the absorbance units (au), into colony forming units (cfu). The transformation factor was derived by plate counting of the starting REL606 strain grown at 37°C in the RTS-1C with the same settings and media of the laboratory evolution experiment. The culture was sampled at mid exponential phase (OD ≈ 0.25) and plated in replicates of six serial dilutions in M9 agar with glucose. The colonies were counted after the overnight incubation at 37°C. With this method, we obtained the following average transformation factor:

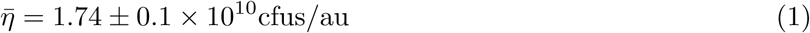

The caveats of using this correction factor as opposed to a calibration curve is that it relies on extrapolation and ignores possible non-linearities when approaching saturation. For our purposes, however, this method is sufficient as our cultures do not spend much time in stationary phase.

The cumulative number of cell divisions (CCD) was also calculated. The CCD provides an alternative measure of a meaningful time-scale for an adaptive evolution experiment and relies on the fact that most mutations occur at the moment of division (Lee et al. 2011). We calculated the number of generations and CCD for all of our populations by splitting the growth data by temperature taking the initial and final optical density at each temperature segment. The following expressions were used to calculate the number of generations and the CCD:

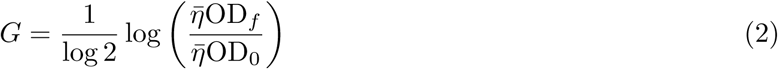

And for all the *m* segments during the adaptive laboratory evolution:

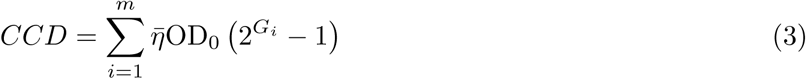

where OD_0_ is the initial OD_830_ optical density and OD_*f*_ is the final OD_830_ optical density of the growth segment considered.

The number of generations and cumulative cell divisions were determined for each experimental population at each temperature (see Table S2). Overall the OD-dependent treatments did not differ significantly from the random treatment in either generations or CCD as expected. They also did not show a significant asymmetry between the experimental temperatures. The populations with a random environment treatment went through slightly more generations due to the finite duration of the 15°C cycle, which in turn could take place during a lag period not amounting to substantial growth. These populations also showed a marked asymmetry between the experimental temperatures in both generations and CCD indicating that they spent more generations in the 43°C. Nevertheless, it is important to note that the amount of generations or cell divisions spent in the 15°C environment is comparable for all treatments. The discrepancies in the total number of generations are explained by two instrumental problems. The first relates to some incomplete data as some of the instruments stopped recording the optical density measurements. The second, as a consequence of instrument malfunction we had to stop some of our populations earlier than others. These populations however did not show significantly fewer mutations suggesting that most of the adaptive mutations occurred in the early stages of the experiments.

### 6.4 Representative clone isolation

From both the starting and final archived heterogeneous populations representative clones were isolated by picking single colonies on M9 glucose agar plates grown at 37°C. The picked clones were grown for a cycle in the experimental conditions. The resulting 20 mL culture was aliquoted in two halves. One half was immediately frozen in glycerol at −80°C for future studies while the other half was pelleted for genome sequencing.

### 6.5 Genome Sequencing and Analysis

For the study of the evolutionary dynamics of our experimental populations, for the final points the genomic sequencing of the populations was conducted by directly pelleting the remaining culture after the transfer to fresh media. In doing so, we follow the recommendation of (Sprouffske et al. 2016) that analyzed the effects of the archiving process on the mutational composition of heterogeneous bacterial populations to avoid bias introduced by selection acting on the freezing-thawing cycle. For the starting points, 30 *μL* of the glycerol stock were sequenced. All sequenced samples were treated as follows. The genomic DNA was isolated with the PureLink genomic DNA mini kit. The extracted genomic DNA was evaluated for quality using Nano-drop absorbance rations and quantified using Qubit dsDNA high sensitivity assay kit. All sequencing was done with 2×75bp paired-end massively parallel sequencing on an Illumina MiSeq (San Diego, USA). Structural and single nucleotide changes at the genomic level were predicted from the Illumina reads using *breseq* (version 0.28.1) assuming that they pertained to an heterogeneous population (Bernstein & Carlson 2014). The reads were aligned to an updated version of the *Escherichia coli* REL606 reference genome file (GenBank accession NC 012967.1). All mutations reported were manually curated to ensure validity.

To better characterize the genomic composition of the starting populations, an additional sequencing round was performed using the NexSeq. The DNA was extracted from 30*μL* of the glycerol stocks using the PureLink genomic DNA mini kit. We fragmented the DNA using 300 bp setting using Covaris. Libraries were prepared using KAPA LT DNA library prep kit with dual size selection to get the library size range between 200-400 bp. Libraries were quantified using Qubit and qualitated by Agilent bioanalyzer. Libraries were further quantified using KAPA QPCR and sequenced on Illumina NEXtseq500 as paired end 2 × 150 bp. The coverage of both populations averaged to 6000. Of all the low frequency mutants detected and not reported in Table S1, none was carried over in the laboratory evolution experiment.

### 6.6 Growth characterization

The representative clones were resurrected at 37°C from the glycerol stocks into M9 media supplemented with glucose 4 *g/L*. After almost reaching saturation, they were acclimated to the tested temperature by transferring 100 *μL* of the culture to 20 mL of fresh M9 media with glucose at the desired temperature. Then, halfway through the exponential phase, the cultures were transferred the same way to triplicates (or sometimes up to six replicates for the strains with greater variance) at the same tested temperature. Each of these experiments was done using the same media batch and the appropriate reference clone (either our starting REL606 or REL607 isolated clones). The growth rate was calculated by fitting a spline to the growth data and calculating the derivative of this fitted curve. The maximum growth rate was then given by the maximum of this derivative. The smoothing parameter was adjusted for the measurement noise but kept constant within each comparison. The relative growth rate was calculated by dividing the average value of the strain measured by the average value of its ancestral reference clone. Significance was assessed using a two sample t-test with directional null hypothesis. All calculations were performed in R version 3.5.1.

To assess the effects of the previous environment on the growth rate, we modified the protocol to have two passages with replicates in the other temperature after acclimation to the target temperature. For example, if the acclimation passage was in 43°C, then we performed two additional passages at 15°C to monitor the previous effects at 43°C on growth at 15°C (see Supplementary Figure 2).

### 6.7 Biolog Assays

Phenotypic assays for each of the 24 clones isolated from the final evolved populations and the two clones isolated from the ancestral populations were performed using Biolog Gen III plates (Biolog, Hayward, CA). These are 96 well plates comprised of 70 metabolic wells with one negative control well and 22 chemical sensitivity wells with one positive control well. Protocols were adapted from the methods in (Cooper & Lenski 2000, Hug & Gaut 2015). The bacteria strains were revived from glycerol stocks and incubated at 37°C for one growth cycle in fresh M9 minimal media supplemented with glucose 4 g/L. This amount of glucose was chosen to avoid starvation cycles. The strains were then transferred to fresh M9 media with glucose and grown again at 37°C. After reaching the middle of exponential growth phase as defined by an OD_830_ of approximately 0.25, the samples were spun down at 4000 rpm for 10 minutes in order to form a cell pellet. The cell pellet was then washed in sterile PBS, and resuspended in 10 mL PBS by vortexing for 30 seconds. The solution was then spun down again at 4000 rpm for 10 minutes, washed with PBS, and resuspended again in 20 mL of PBS. The resulting suspensions were aliquoted into 10*mL* vials of Biolog Inoculating Fluid A. Bacteria were aliquoted in the inoculating fluid with the appropriate volume to obtain 95% transmittance, which was verified by spectrophotometry to within 3%. The equation *OD* = −*log*_10_(*T)*, where *OD* is the *OD*_830_ and *T* is the transmittance, was used to calculate the dilution of the bacteria into the inoculating fluid.

After being aliquoted into the inoculating fluid, the bacteria and inoculating fluid mixture was vortexed for 20 seconds to ensure mixing, and then 100*μ*L of inoculating fluid bacteria mixture was aliquoted into each of the 96 wells of the Biolog Gen III plates. After being plated, the plates were labeled and placed in incubators at the appropriate temperatures, either 15°C, 37°C, or 43°C, and allowed to grow. Plates grown at 37°C were grown for 24 - 36 hours, plates grown at 43 °C were grown for 48 - 60 hours, and plates grown at 15°C were grown for 72 - 96 hours to ensure saturation of bacterial growth. After saturation, the plates were measured on a BioTek plate reader (BioTek, Winooski, VT) at 590 nm. The measurements were repeated over time to ensure the stability of the reading.

All Biolog plates were performed in technical triplicates for each of the 24 evolved clones at each temperature, for a total of 24 × 3 × 3 = 216 plates. The two ancestral strains were performed in six technical replicates for each of the assayed temperatures in order to better assess the ancestral phenotypic response, for a total of 2 × 6 × 3 = 36 ancestor plates.

### 6.8 Analysis of Biolog Data

The biolog optical density measurements (OD_590_) were processed following the instructions of the maker. First, normalization of each plate was performed by subtracting its negative control from all metabolic wells. Additionally, the chemical sensitivity assays were normalized for each plate by subtracting those wells from the value of the positive control. Both the negative and positive controls were removed from the analysis. Wells 86 and 94, which contain two different tetrazolium dyes (which can be interpreted as two additional positive growing conditions as no inhibitory effect from the dies was observed in any of the populations) had consistently higher values than the rest of the wells for all the evolved and ancestral samples but the values did not vary between the samples. Thus, these two wells were removed from our analysis. This left us with 92 phenotypic tests for each sample, represented as a vector in 92-dimensional space. An alternative data processing was also performed following (Hug & Gaut 2015). The conclusions were robust to both data processing methods. Additionally for a couple of samples, data quality control was done to correct for any technical replicates with some inconsistent well values, and thus were corrected by imputation by taking the mean well value of the remaining replicates. The resulting processed data was then centered. Clustering of the output of the Biolog data was performed using Ward’s algorithm for hierarchical clustering with Euclidean distance.

In order to assess the direction of evolved strains’ phenotypic responses with respect to their respective ancestor, we followed the methodology put forward by (Hug & Gaut 2015). Because of the different mutational backgrounds of the ancestral strains, we performed the analysis separately for the REL606 and REL607 lineages. The dimensionality reduction of the phenotypic measurements was performed using principal component analysis (PCA) on the processed data to remove correlated variables due to the Biolog plate design. The centroids of each set of ancestral replicates at the three different temperatures were calculated by k-means clustering. For each lineage, the first four principal components were selected according to the criterion in (Peres-Neto et al. 2005) amounting to ∼70% of the variance. Additional criteria to select informative principal components like the talus plot (Henningsson et al. 2018) were also taken into consideration obtaining similar results. T-test hypothesis testing with family-wise error rate correction was conducted to determine the direction of the phenotypic response for each evolved strain and retained principal component in relation to the ancestral response at the optimal and the stressed temperatures. Following (Hug & Gaut 2015), the responses were classified as restorative, partially restorative, reinforced, unrestored, novel, or uninformative. All calculations were performed in R version 3.5.3, including the mutational fingerprint (Gu et al. 2016), and Python 2.7.15.

## Supporting Information

### Supplementary Figures and Tables

**Figure S 1:**
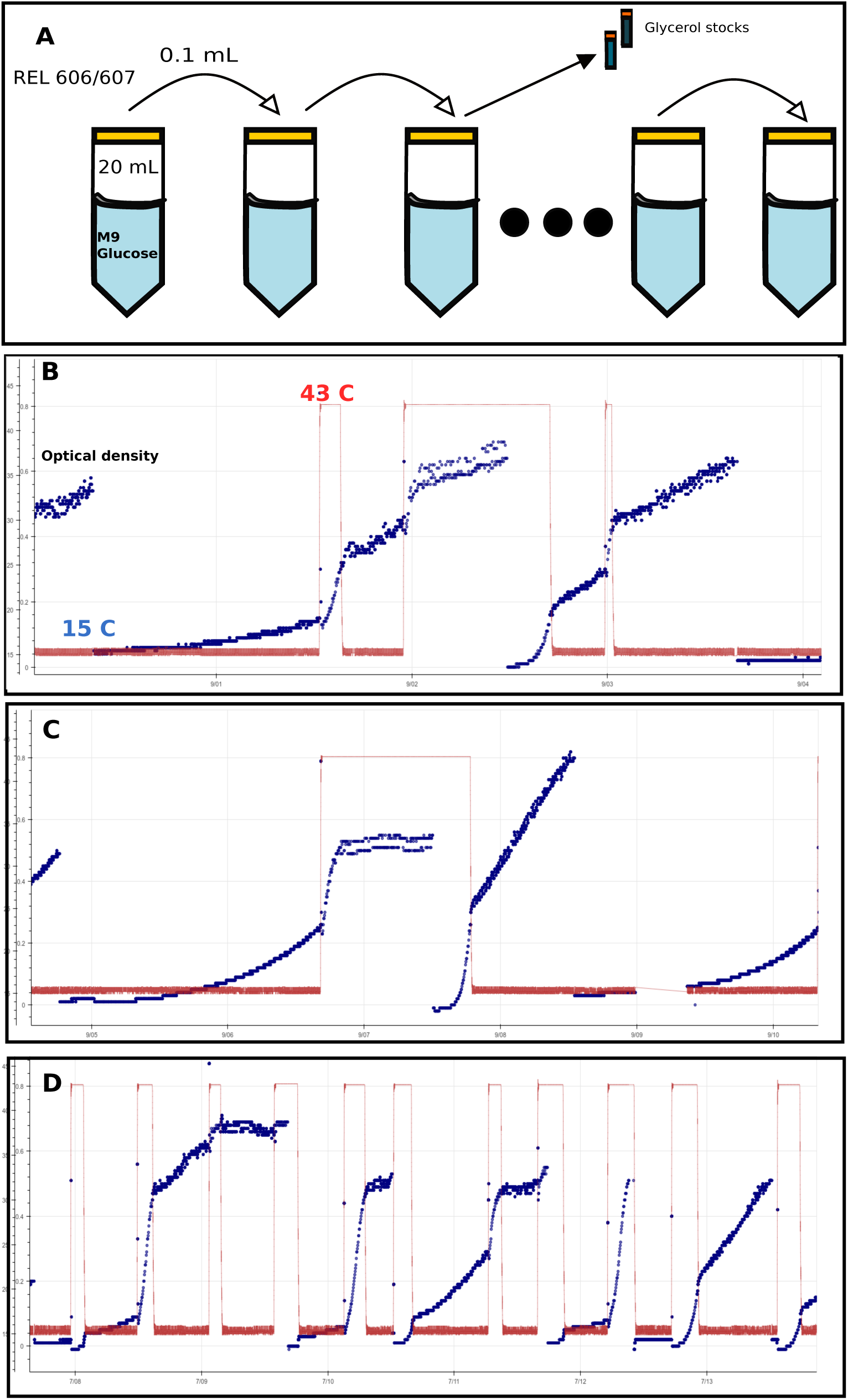
Experimental evolution design of the sample optical density and temperature trajectories for each of the three treatments. **A**: The transfer schema performed during the laboratory evolution experiment. **B**: A sample trajectory from the OD-dependent fast treatment.**C**: A sample trajectory from the OD-dependent slow treatment.**D**: A sample trajectory from the random treatment. Red lines show temperature in Celsius while blue dots represent OD_830_ measurements.

**Table S 1:** Background mutations found in our starting points of our adaptive laboratory experiment.

| Gene | 606 | 607 | Function | Mutation | Protein Change |
| --- | --- | --- | --- | --- | --- |
| araA | 0 | 100 | Larabinose isomerase | SNP T/C | D92G (GAC→GGC) |
| recD | 0 | 100 | exonuclease V (RecBCD complex) | SNP T/C | V10A (GTT→GCT) |
| nadR | 100 | 0 | nicotinamidenucleotide adenylyltransferase | Coding +G at 4,616,183:1 | Coding (655/1233 nt) |
| nadR | 0 | 71.3 | nicotinamidenucleotide adenylyltransferase | SNP G/T | E367* stop |
| infB | 100 | 0 | translation initiation factor IF2 | SNP A/T | V85D |
| rbsD-[rbsR] | 100 | 0 | multiple genes | IS150 mediated $\Delta$ 5,874bp at 3894997 | Deleted rbsD, rbsA, rbsC, rbsB, rbsK, [rbsR] |
| yhjY/tag | 0 | 20 | hypothetical protein/DNA glycosylase | Intergenic G/A | Intergenic (-42/-116) |
| uxaB | 19.5 | 0 | tagaturonate reductase | SNP C/A | R429L |
| cybB-[ydcG] | 75.4 | 0 | multiple genes | IS1 mediated $\Delta$ 6,496bp at 1460927 | Deleted cybB, ydcA, hokB, mokB, trg, ydcI, ydcJ, [ydcG] |
| ECB.02621 | 0 | 76.9 | conserved hypothetical protein | $\Delta$ 122 bp at 2,792,849 | Deletion |

**Table S 2:** Generations and cumulative cell divisions of our final populations for each of the three environmental treatments, where F-, S-, and R-606/607-replicate# stands for fast, slow, and random fluctuating environmental treatments, respectively.

| Strain | Date stopped | 43 C° CCD (x10<sup>10</sup>) | 15 C° CCD (x10<sup>10</sup>) | Total CCD | 43 C° Generations | 15 C° Generations | Total Generations |
| --- | --- | --- | --- | --- | --- | --- | --- |
| F606-1 | 09/13/16 | 27.52 | 26.51 | 54.03 | 254.44 | 230.05 | 484.49 |
| F606-2 | 09/07/16 | 46.1 | 41.56 | 87.67 | 358.57 | 348.2 | 706.77 |
| F606-3 | 09/08/16 | 49.73 | 66.05 | 115.78 | 307.57 | 264.52 | 572.09 |
| F606-4 | 07/16/16 | 18.29 | 19.72 | 38.01 | 160.99 | 151.89 | 312.88 |
| F607-1 | 09/11/16 | 18.02 | 19.28 | 37.3 | 166.37 | 172.71 | 339.08 |
| F607-2 | 09/12/16 | 34.92 | 31.03 | 65.95 | 297.71 | 285.08 | 582.64 |
| F607-3 | 09/10/16 | 41.75 | 25.25 | 67 | 328.96 | 214.24 | 543.2 |
| F607-4 | 07/17/16 | 17.42 | 21.17 | 38.59 | 168.49 | 174.18 | 342.67 |
| S606-1 | 09/07/16 | 39.75 | 38.18 | 77.92 | 337.53 | 328.86 | 666.39 |
| S606-2 | 07/23/16 | 29.25 | 28.1 | 57.35 | 255.09 | 240.76 | 495.85 |
| S606-3 | 09/09/16 | 29.44 | 27.08 | 56.52 | 257.37 | 250.57 | 507.93 |
| S606-4 | 08/14/16 | 31.91 | 37.59 | 69.5 | 288.2 | 300.75 | 588.94 |
| S607-1 | 09/09/16 | 38.22 | 40.26 | 78.48 | 328.65 | 331.3 | 659.95 |
| S607-2 | 08/07/16 | 25.37 | 26.25 | 51.62 | 218.85 | 224.82 | 443.67 |
| S607-3 | 09/09/16 | 40.04 | 35.95 | 75.99 | 359.64 | 301.8 | 661.44 |
| S607-4 | 09/14/16 | 120.98 | 43.47 | 164.45 | 357.18 | 335.27 | 692.45 |
| R606-1 | 09/09/16 | 50.63 | 26.36 | 76.99 | 419.3 | 327.41 | 746.71 |
| R606-2 | 09/09/16 | 148.66 | 32.91 | 181.57 | 346.4 | 444.14 | 790.54 |
| R606-3 | 09/13/16 | 50.96 | 24.77 | 75.74 | 368.55 | 330.77 | 699.32 |
| R606-4 | 09/08/16 | 50.06 | 27.26 | 77.32 | 496.08 | 185.65 | 681.74 |
| R607-1 | 09/09/16 | 63.87 | 32.64 | 96.51 | 464.3 | 368.02 | 832.31 |
| R607-2 | 09/09/16 | 170.96 | 28.39 | 199.35 | 442.64 | 315.12 | 757.76 |
| R607-3 | 09/09/16 | 53.54 | 31.86 | 85.4 | 581.39 | 170.02 | 751.41 |
| R607-4 | 09/09/16 | 49.4 | 30.61 | 80.01 | 551.35 | 185.42 | 736.87 |

**Table S3A 0:** Mutations found in our final REL606-derived populations evolved in the random environmental treatment. Only the mutations that are present in more than 5% of the populations are shown.

| Gene | R606-1 | R606-2 | R606-3 | R606-4 | Mutation Annotation |
| --- | --- | --- | --- | --- | --- |
| ***Ancestral*** |  |  |  |  |  |
| rbsD-[rbsR] | 100 | 100 | 100 | 100 | IS150-mediated $\Delta$ 5,874 bp at 3,894,997 |
| cybB-[ydcG] | 100 | 100 | 0 | 100 | IS1-mediated Δ6,496 bp at 1,460,928 |
| infB | 100 | 100 | 100 | 100 | V85D (G<u>T</u>C→G<u>A</u>C) |
| nadR | 100 | 100 | 100 | 100 | Coding (658/1233 nt) +G at 4,616,186 |
| ***Heat Shock:*** |  |  |  |  |  |
| insL2/lon | 72.4 | 0 | 0 | 0 | Intergenic IS mediated Δ1,350 bp at 429,504 +24 bp at 430,879 |
| insL2/lon | 0 | 100 | 0 | 100 | Intergenic (-92/-64) *G* → *T* at 430,879 |
| insL2-[ppiD] | 0 | 0 | 48.9 | 0 | Amplification at [430,618-434,631] |
| ***Cold Shock:*** |  |  |  |  |  |
| deaD | 60.2 | 0 | 0 | 0 | Coding $\Delta$ 142 bp at 3,241,224 |
| ***Membrane and Cell Wall:*** |  |  |  |  |  |
| mrdA | 52.3 | 0 | 0 | 0 | D312A (GAC→GCC) |
| mrdA | 0 | 18.6/6.1 | 0 | 0 | V175I (G<u>T</u>C→<u>A</u>TC)/D354A (G<u>A</u>C→G<u>C</u>C) |
| mrdB | 0 | 65.2 | 0 | 0 | S37N (A<u>G</u>C→ A<u>A</u>C) |
| mreC | 33.3 | 0 | 0 | 0 | Coding (1077/1104 nt) +C at 332,679 |
| mreC | 0 | 0 | 0 | 9.5/7.5 | S141C (A<u>G</u>C→T<u>G</u>C)/R113S (C<u>G</u>T→A<u>G</u>T) |
| mreD | 0 | 0 | 0 | 35/17 | P28L (C<u>C</u>G→C<u>T</u>G)/ P36L |
| ***Transporter:*** |  |  |  |  |  |
| sapA | 0 | 59.1 | 0 | 0 | Y187N (TAT→AAT) |
| pstA | 0 | 12.3 | 0 | 0 | W264* (T<u>G</u>G→T<u>G</u>A) |
| ***Metabolic:*** |  |  |  |  |  |
| waaL | 0 | 0 | 0 | 23.5 | G101R |
| fadH | 0 | 0 | 6.1 | 0 | L216L |
| napA | 0 | 0 | 100 | 0 | Coding (1510/2487 nt) Δ1bp at 2,251,232 |
| edd/zwf | 0 | 0 | 100 | 0 | Intergenic (-30/+205) A→C at 1,913,210 |
| ***Others:*** |  |  |  |  |  |
| gyrB | 0 | 0 | 100 | 0 | R732H (C<u>G</u>C→C<u>A</u>C) |
| thiC/rsd | 0 | 0 | 100 | 0 | Intergenic (-166/+68) A→T at 4,175,876 |
| ogrK-[ECB 02013] | 0 | 18.5 | 0 | 0 | $\Delta$ 2,2146 bp at 2,100,308 |
| proQ | 0 | 0 | 100 | 0 | Coding (333-345/699 nt) $\Delta$ 13bp at 1,893,767 |
| insC/ECB_01906 | 8.3 | 0 | 0 | 0 | $\Delta$ 7bp at 2,001,021 intergenic (641/+8) |

**Table S3B 0:** Mutations found in our final REL607-derived populations evolved in the random environmental treatment. Only the mutations that are present in more than 5% of the populations are shown.

| Gene | R607-1 | R607-2 | R607-3 | R607-4 | Mutation Annotation |
| --- | --- | --- | --- | --- | --- |
| ***Ancestral:*** |  |  |  |  |  |
| araA | 100 | 100 | 100 | 100 | D92G (G<u>A</u>C→G<u>G</u>C) |
| recD | 100 | 100 | 100 | 100 | V10A (G<u>T</u>T→G<u>C</u>T) |
| yhjY/tag | 100 | 38.0 | 0 | 100 | Intergenic (-42/-116) G→A at 3,643,434 |
| nadR | 0 | 65.1 | 100 | 0 | E367* stop (G<u>A</u>G→<u>T</u>AG) |
| ECB.02621 | 0 | 66.7 | 100 | 0 | Coding Δ 122 bp at 2,792,849 |
| ***Regulatory:*** |  |  |  |  |  |
| rho | 0 | 100 | 100 | 100 | I15N (A<u>T</u>C→A<u>A</u>C) |
| rho | 55.4/34.6 | 0 | 0 | 0 | R221C (<u>C</u>GT→<u>T</u>GT) /H295Y (<u>C</u>AT→<u>T</u>AT) |
| ***Heat Shock:*** |  |  |  |  |  |
| lon | 0 | 13 | 0 | 0 | E4E (G<u>A</u>G→G<u>A</u><u>A</u>) |
| insL2/lon | 100 | 0 | 0 | 0 | Intergenic (-92/-64) G → T at 430,879 |
| insL2/lon | 0 | 0 | 100 | 100 | Intergenic (-48/-108) C → T at 430,835 |
| ***Cold Shock:*** |  |  |  |  |  |
| deaD | 0 | 53.3/7.4 | 0 | 0 | D321A (G<u>A</u>T→G<u>C</u>T) /L40Q (C<u>T</u>G→C<u>A</u>G) |
| deaD | 0 | 0 | 39.5 | 0 | Coding (1223/1890 nt) +66bp at 3,241,872 |
| ***Membrane and Cell Wall:*** |  |  |  |  |  |
| dacA | 100 | 0 | 0 | 0 | K76T (A<u>A</u>A→A<u>C</u>A) |
| mrdA | 0 | 0 | 0 | 100 | Y397S (T<u>A</u>C→T<u>C</u>C) |
| mreC | 0 | 0 | 8.8 | 0 | L84Q (C<u>T</u>G→C<u>A</u>G) |
| yjiY/hpaC | 0 | 0 | 94.2 | 0 | Intergenic (-347/+46) T/A at 456,960 |
| ***Metabolic:*** |  |  |  |  |  |
| icdA | 0 | 5.8 | 0 | 0 | D158D G<u>A</u>C→G<u>A</u><u>T</u>) |
| nadR | 100 | 0 | 0 | 100 | Coding (165/1233 nt) Δ1bp at 4,615,693 |
| ***Others:*** |  |  |  |  |  |
| yeaP/yeaQ | 0 | 0 | 0 | 5.5 | Intergenic (+34/+233) T/C at 1,856,003 |

**Table S3C 0:** Mutations found in our final REL6060-derived populations evolved in the fast deterministic environmental treatment. Only the mutations that are present in more than 5% of the populations are shown.

| Gene | F606-1 | F606-2 | F606-3 | F606-4 | Mutation |
| --- | --- | --- | --- | --- | --- |
| ***Ancestral*** |  |  |  |  |  |
| rbsD-[rbsR] | 100 | 100 | 100 | 100 | IS150-mediated $\Delta$ 5,874 bp at 3,894,997 |
| cybB-[ydcG] | 29.8 | 100 | 100 | 100 | IS1-mediated Δ6,496 bp at 1,460,928 |
| infB | 100 | 100 | 100 | 100 | V85D (G<u>T</u>C→G<u>A</u>C) |
| nadR | 100 | 100 | 100 | 100 | Coding (658/1233 nt) +G at 4,616,186 |
| uxaB | 61.4 | 0 | 0 | 0 | R429L (C<u>G</u>T→C<u>T</u>T) |
| ***Regulatory:*** |  |  |  |  |  |
| ycjW | 12.5 | 0 | 0 | 0 | IS150 (-) +3 bp coding (177-179/999 nt) |
| elaD | 0 | 0 | 0 | 100 | $\Delta$ 1 bp at 2,329,952 coding (734/1212 nt) |
| ***Heat Shock:*** |  |  |  |  |  |
| yjeH/groES | 0 | 100 | 0 | 0 | Δ3 bp at 4,349,333 intergenic (-200/-74) |
| [insL-2]-[ppiD] | 0 | 0 | 49.7 | 0 | Amplification at [430689-435601] |
| [dsbD]-efp/ecnA | 0 | 0 | 0 | 62.7 | Amplification at [4343517-4354989] |
| lon | 0 | 0 | 6.7 | 0 | D445E |
| insL2/lon | 8.3 | 0 | 0 | 0 | Intergenic (-24/-132) G→A at 430,811 |
| insL2/lon | 71.3 | 0 | 0 | 0 | Intergenic (-48/-108) C → T at 430,835 |
| insL2/lon | 0 | 94.0 | 0 | 94.3 | Δ1350 bp at 429505 non-coding and intergenic (-68/-88) |
| ***Metabolic:*** |  |  |  |  |  |
| hemF | 0 | 51.3 | 0 | 0 | Δ27bp at 2,481,400 coding (657-683/900 nt) |
| dsdA | 6.1 | 0 | 0 | 0 | I265T (A<u>T</u>C→A<u>C</u>C) |
| metA | 13.7 | 0 | 0 | 0 | I124L (<u>A</u>TC→<u>C</u>TC) |
| ***Membrane and Cell Wall:*** |  |  |  |  |  |
| mrdA | 0 | 0 | 61.5 | 0 | D354A (G<u>A</u>C→G<u>C</u>C) |
| mrdA | 0 | 0 | 14.3 | 0 | N428D (<u>A</u>AC→<u>G</u>AC) |
| mrdB | 57.6 | 0 | 0 | 0 | R208S (<u>C</u>GC→<u>A</u>GC) |
| mreD | 0 | 0 | 5.8 | 0 | V161A (G<u>T</u>G→G<u>C</u>G) |
| ***Others:*** |  |  |  |  |  |
| rpmH | 6 | 0 | 0 | 0 | Amplification +CG at 3,844,715 coding |

**Table S3D 0:**
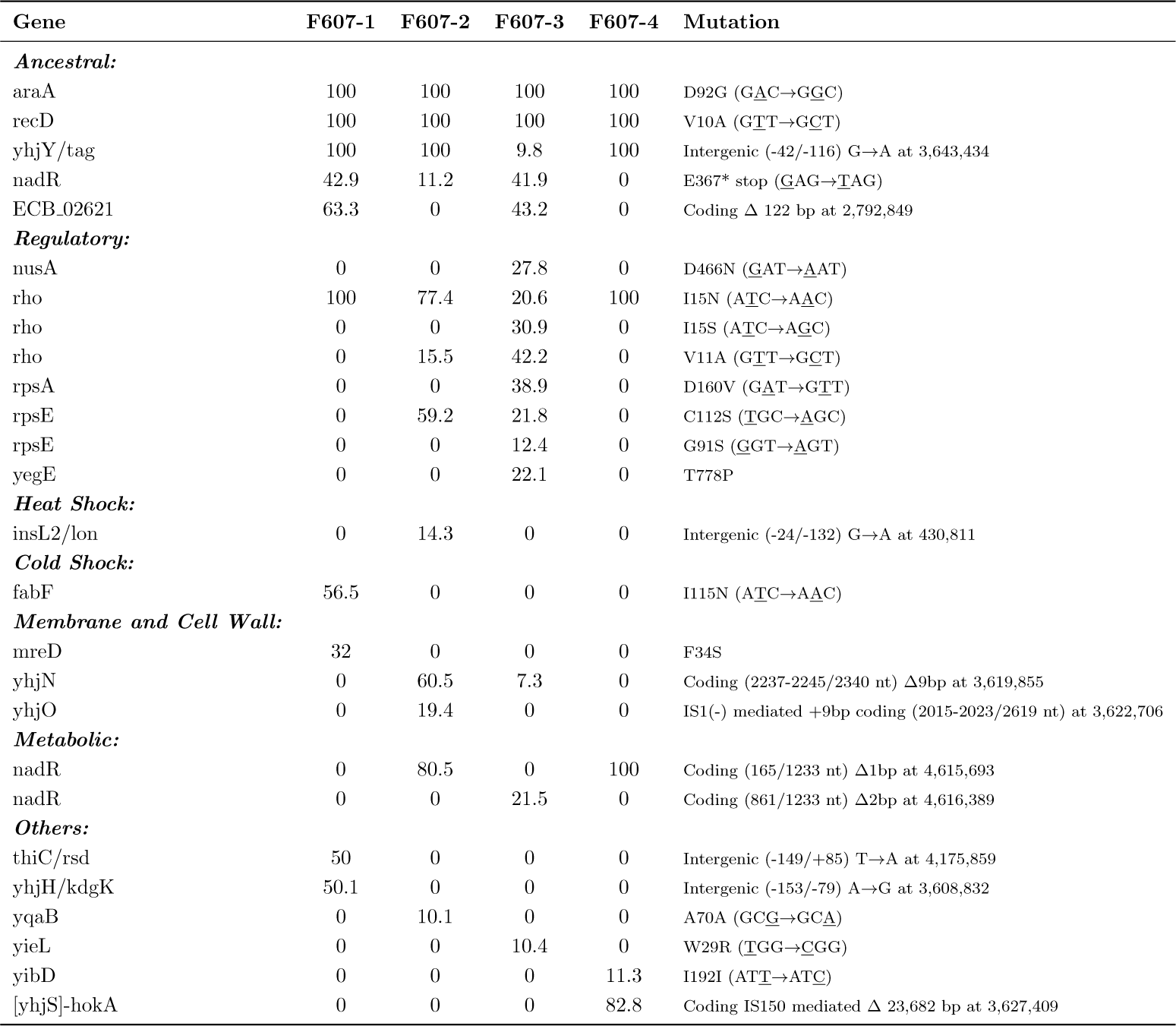
Mutations found in our final REL607-derived populations evolved in the fast deterministic environmental treatment. Only the mutations that are present in more than 5% of the populations are shown.

| Gene | F607-1 | F607-2 | F607-3 | F607-4 | Mutation |
| --- | --- | --- | --- | --- | --- |
| *Ancestral:* |  |  |  |  |  |
| araA | 100 | 100 | 100 | 100 | D92G (G<u>A</u>C→G<u>G</u>C) |
| recD | 100 | 100 | 100 | 100 | V10A (G<u>T</u>T→G<u>C</u>T) |
| yhjY/tag | 100 | 100 | 9.8 | 100 | Intergenic (-42/-116) G→A at 3,643,434 |
| nadR | 42.9 | 11.2 | 41.9 | 0 | E367* stop (G<u>A</u>G→<u>T</u>AG) |
| ECB_02621 | 63.3 | 0 | 43.2 | 0 | Coding $\Delta$ 122 bp at 2,792,849 |
| *Regulatory:* |  |  |  |  |  |
| nusA | 0 | 0 | 27.8 | 0 | D466N (G<u>A</u>T→<u>A</u>AT) |
| rho | 100 | 77.4 | 20.6 | 100 | I15N (A<u>T</u>C→A<u>A</u>C) |
| rho | 0 | 0 | 30.9 | 0 | I15S (A<u>T</u>C→A<u>G</u>C) |
| rho | 0 | 15.5 | 42.2 | 0 | V11A (G<u>T</u>T→G<u>C</u>T) |
| rpsA | 0 | 0 | 38.9 | 0 | D160V (G<u>A</u>T→G<u>T</u>T) |
| rpsE | 0 | 59.2 | 21.8 | 0 | C112S (T<u>G</u>C→<u>A</u>GC) |
| rpsE | 0 | 0 | 12.4 | 0 | G91S (G<u>G</u>T→<u>A</u>GT) |
| yegE | 0 | 0 | 22.1 | 0 | T778P |
| *Heat Shock:* |  |  |  |  |  |
| insL2/lon | 0 | 14.3 | 0 | 0 | Intergenic (-24/-132) G→A at 430,811 |
| *Cold Shock:* |  |  |  |  |  |
| fabF | 56.5 | 0 | 0 | 0 | I115N (A<u>T</u>C→A<u>A</u>C) |
| *Membrane and Cell Wall:* |  |  |  |  |  |
| mreD | 32 | 0 | 0 | 0 | F34S |
| yhjN | 0 | 60.5 | 7.3 | 0 | Coding (2237-2245/2340 nt) Δ9bp at 3,619,855 |
| yhjO | 0 | 19.4 | 0 | 0 | IS1(-) mediated +9bp coding (2015-2023/2619 nt) at 3,622,706 |
| *Metabolic:* |  |  |  |  |  |
| nadR | 0 | 80.5 | 0 | 100 | Coding (165/1233 nt) Δ1bp at 4,615,693 |
| nadR | 0 | 0 | 21.5 | 0 | Coding (861/1233 nt) Δ2bp at 4,616,389 |
| *Others:* |  |  |  |  |  |
| thiC/rsd | 50 | 0 | 0 | 0 | Intergenic (-149/+85) T→A at 4,175,859 |
| yhjH/kdgK | 50.1 | 0 | 0 | 0 | Intergenic (-153/-79) A→G at 3,608,832 |
| yqaB | 0 | 10.1 | 0 | 0 | A70A (G<u>C</u>G→G<u>C</u>A) |
| yieL | 0 | 0 | 10.4 | 0 | W29R (T<u>G</u>G→<u>C</u>GG) |
| yibD | 0 | 0 | 0 | 11.3 | I192I (A<u>T</u>T→A<u>T</u>C) |
| [yhjS]-hokA | 0 | 0 | 0 | 82.8 | Coding IS150 mediated $\Delta$ 23,682 bp at 3,627,409 |

**Table S3E 0:**
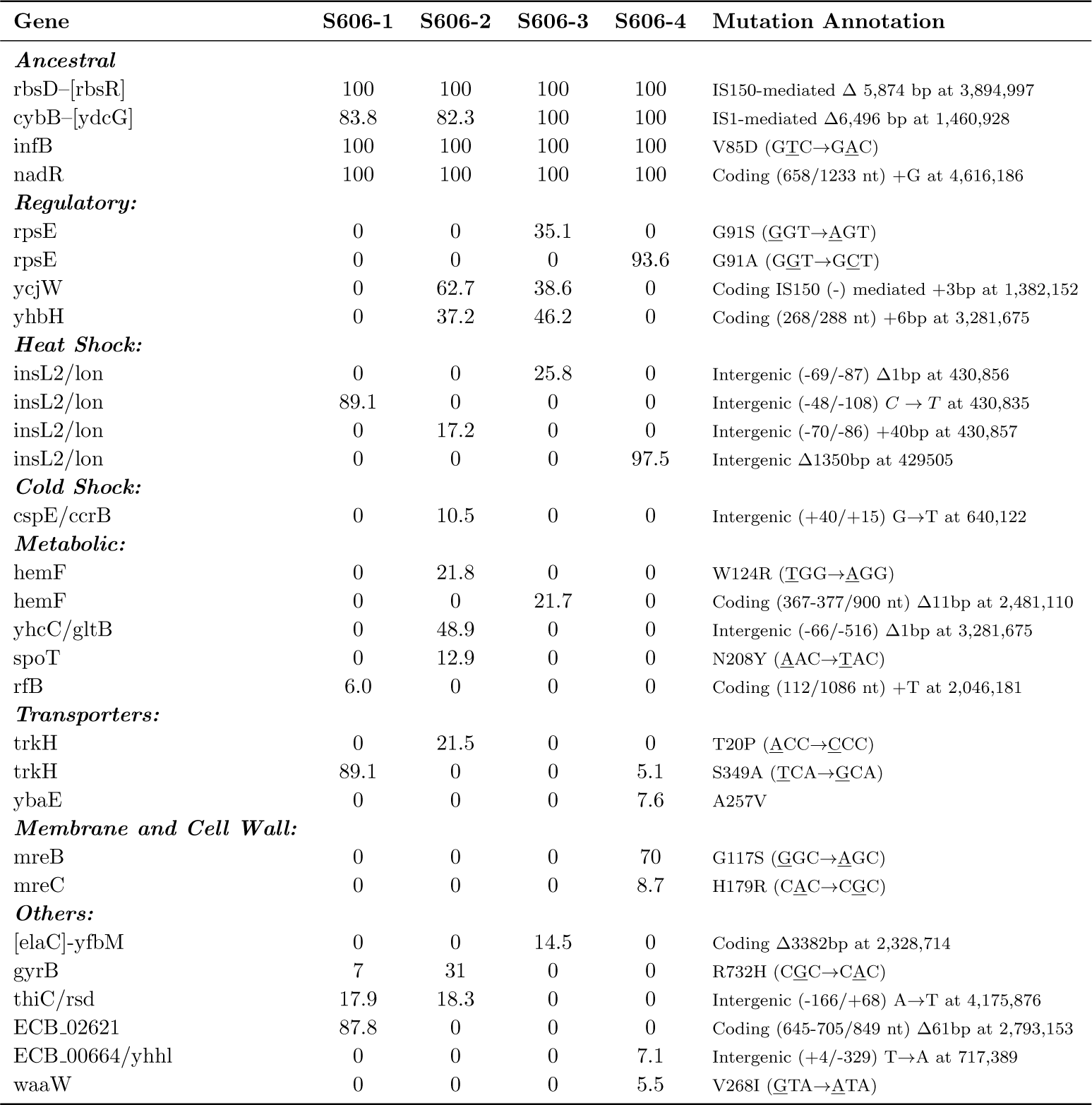
Mutations found in our final REL606 derived populations evolved in the slow deterministic environment. Only the mutations that are present in more than 5% in the populations are shown.

| Gene | S606-1 | S606-2 | S606-3 | S606-4 | Mutation Annotation |
| --- | --- | --- | --- | --- | --- |
| ***Ancestral*** |  |  |  |  |  |
| rbsD-[rbsR] | 100 | 100 | 100 | 100 | IS150-mediated $\Delta$ 5,874 bp at 3,894,997 |
| cybB-[ydcG] | 83.8 | 82.3 | 100 | 100 | IS1-mediated Δ6,496 bp at 1,460,928 |
| infB | 100 | 100 | 100 | 100 | V85D (<u>G</u>T<u>C</u>→<u>G</u>A<u>C</u>) |
| nadR | 100 | 100 | 100 | 100 | Coding (658/1233 nt) +G at 4,616,186 |
| ***Regulatory:*** |  |  |  |  |  |
| rpsE | 0 | 0 | 35.1 | 0 | G91S (<u>G</u>G<u>T</u>→<u>A</u>G<u>T</u>) |
| rpsE | 0 | 0 | 0 | 93.6 | G91A (<u>G</u>G<u>T</u>→<u>G</u>C<u>T</u>) |
| ycjW | 0 | 62.7 | 38.6 | 0 | Coding IS150 (-) mediated +3bp at 1,382,152 |
| yhbH | 0 | 37.2 | 46.2 | 0 | Coding (268/288 nt) +6bp at 3,281,675 |
| ***Heat Shock:*** |  |  |  |  |  |
| insL2/lon | 0 | 0 | 25.8 | 0 | Intergenic (-69/-87) Δ1bp at 430,856 |
| insL2/lon | 89.1 | 0 | 0 | 0 | Intergenic (-48/-108) *C* → *T* at 430,835 |
| insL2/lon | 0 | 17.2 | 0 | 0 | Intergenic (-70/-86) +40bp at 430,857 |
| insL2/lon | 0 | 0 | 0 | 97.5 | Intergenic Δ1350bp at 429505 |
| ***Cold Shock:*** |  |  |  |  |  |
| cspE/ccrB | 0 | 10.5 | 0 | 0 | Intergenic (+40/+15) G→T at 640,122 |
| ***Metabolic:*** |  |  |  |  |  |
| hemF | 0 | 21.8 | 0 | 0 | W124R (<u>T</u>G<u>G</u>→<u>A</u>G<u>G</u>) |
| hemF | 0 | 0 | 21.7 | 0 | Coding (367-377/900 nt) Δ11bp at 2,481,110 |
| yhcC/gltB | 0 | 48.9 | 0 | 0 | Intergenic (-66/-516) Δ1bp at 3,281,675 |
| spoT | 0 | 12.9 | 0 | 0 | N208Y (<u>A</u>A<u>C</u>→<u>T</u>A<u>C</u>) |
| rfB | 6.0 | 0 | 0 | 0 | Coding (112/1086 nt) +T at 2,046,181 |
| ***Transporters:*** |  |  |  |  |  |
| trkH | 0 | 21.5 | 0 | 0 | T20P (<u>A</u>C<u>C</u>→<u>C</u>C<u>C</u>) |
| trkH | 89.1 | 0 | 0 | 5.1 | S349A (<u>T</u>C<u>A</u>→<u>G</u>C<u>A</u>) |
| ybaE | 0 | 0 | 0 | 7.6 | A257V |
| ***Membrane and Cell Wall:*** |  |  |  |  |  |
| mreB | 0 | 0 | 0 | 70 | G117S (<u>G</u>G<u>C</u>→<u>A</u>G<u>C</u>) |
| mreC | 0 | 0 | 0 | 8.7 | H179R (<u>C</u>A<u>C</u>→<u>C</u>G<u>C</u>) |
| ***Others:*** |  |  |  |  |  |
| [elaC]-yfbM | 0 | 0 | 14.5 | 0 | Coding Δ3382bp at 2,328,714 |
| gyrB | 7 | 31 | 0 | 0 | R732H (<u>C</u>G<u>C</u>→<u>C</u>A<u>C</u>) |
| thiC/rsd | 17.9 | 18.3 | 0 | 0 | Intergenic (-166/+68) A→T at 4,175,876 |
| ECB_02621 | 87.8 | 0 | 0 | 0 | Coding (645-705/849 nt) Δ61bp at 2,793,153 |
| ECB_00664/yhhl | 0 | 0 | 0 | 7.1 | Intergenic (+4/-329) T→A at 717,389 |
| waaW | 0 | 0 | 0 | 5.5 | V268I (<u>G</u>T<u>A</u>→<u>A</u>T<u>A</u>) |

**Table S3F 0:** Mutations found in our final REL607-derived populations evolved in the slow deterministic environmental treatment. Only the mutations that are present in more than 5% of the populations are shown.

| Gene | S607-1 | S607-2 | S607-3 | S607-4 | Mutation |
| --- | --- | --- | --- | --- | --- |
| ***Ancestral:*** |  |  |  |  |  |
| araA | 100 | 100 | 100 | 100 | D92G (<u>G</u><u>A</u>C→<u>G</u><u>G</u>C) |
| recD | 100 | 100 | 100 | 100 | V10A (<u>G</u><u>T</u>T→<u>G</u><u>C</u>T) |
| yhjY/tag | 100 | 0 | 100 | 52.6 | Intergenic (-42/-116) G→A at 3,643,434 |
| nadR | 0 | 0 | 100 | 55.5 | E367* stop (<u>G</u>AG→<u>T</u>AG) |
| ECB.02621 | 0 | 100 | 0 | 63.3 | Coding Δ 122 bp at 2,792,849 |
| ***Regulatory:*** |  |  |  |  |  |
| rho | 0 | 0 | 94.8 | 59.1 | I15N (<u>A</u><u>T</u>C→<u>A</u><u>A</u>C) |
| rho | 0 | 0 | 0 | 39.4 | V11A (<u>G</u><u>T</u>T→<u>G</u><u>C</u>T) |
| rpoB | 0 | 0 | 0 | 58.2 | G907S (<u>G</u>GT→<u>A</u>GT) |
| rpsE | 0 | 0 | 65 | 44.3 | G91C (<u>G</u>GT→<u>T</u>GT) |
| nusA | 46.6 | 0 | 0 | 0 | T72N (<u>A</u>CC→<u>A</u>AC) |
| nusA | 8.0 | 0 | 0 | 0 | T72A (<u>A</u>CC→<u>G</u>CC) |
| nusA | 30.0 | 0 | 0 | 0 | W62R (<u>T</u>GG→<u>C</u>GG) |
| stpA | 0 | 0 | 8.9 | 0 | Coding (246-248/405 nt) Δ 3 bp at 2,693,001 |
| yegE | 0 | 0 | 0 | 48.7 | Coding (1456-1462/3306 nt) Δ 7 bp at 2,077,927 |
| ***Heat Shock:*** |  |  |  |  |  |
| insL2/lon | 0 | 0 | 23.8 | 0 | Intergenic (-24/-32) G→A at 430,811 |
| ***Cold Shock:*** |  |  |  |  |  |
| proQ | 0 | 0 | 0 | 12.2 | Coding (364-370/699 nt) Δ 7 bp at 1,893,742 |
| [fabF]-[pabC] | 0 | 32.5 | 0 | 0 | Coding Δ 650 bp at 1,167,222 |
| ***Metabolic:*** |  |  |  |  |  |
| spoT | 0 | 0 | 52.1 | 0 | R701L (<u>C</u>GA→<u>C</u>TA) |
| spoT | 0 | 0 | 0 | 9 | R487H (<u>C</u>GT→<u>C</u>AT) |
| araB | 0 | 0 | 8.2 | 0 | Coding (559/1701 nt) Δ 1bp at 72,294 |
| nadR | 100 | 0 | 100 | 0 | Coding (165/1233 nt) Δ1bp at 4,615,693 |
| ***Membrane and Cell Wall:*** |  |  |  |  |  |
| mrdA | 0 | 0 | 0 | 43.5 | D354A (<u>G</u>AC→<u>G</u>CC) |
| ***Transport:*** |  |  |  |  |  |
| nmpC | 0 | 0 | 0 | 47.1 | Pseudogene (421/606 nt) T→G at 547,858 |
| ***Others:*** |  |  |  |  |  |
| cadC/pheU-[yjeM] | 58 | 0 | 0 | 0 | Amplification at [4,341,178-4,362,612] |
| [fxsA]-[yjeP] | 0 | 71.6 | 0 | 0 | Amplification at [4,347,401-4,366,500] |
| ECB.01992 | 0 | 100 | 0 | 0 | Coding (154/216 nt) +CAGC at 2,103,887 |
| thiC/rsd | 56.5 | 0 | 0 | 0 | Intergenic (-134/+100) A→C at 4,175,844 |
| waaW | 10.1 | 0 | 0 | 0 | Coding (626-631/984 nt) Δ 6 bp at 3,738,225 |

**Table S 4:**
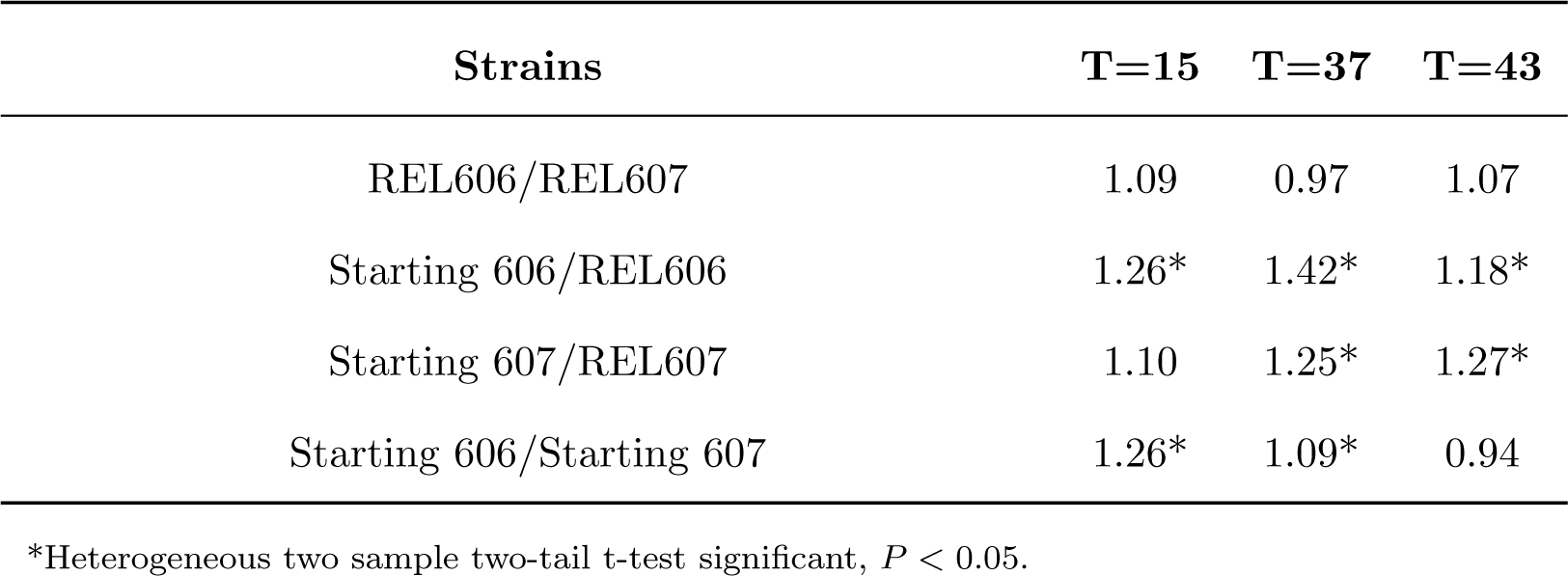
Relative growth of our starting strains, termed starting 606/607, and the original Lenski strains, termed REL606/607.

| Strains | T=15 | T=37 | T=43 |
| --- | --- | --- | --- |
| REL606/REL607 | 1.09 | 0.97 | 1.07 |
| Starting 606/REL606 | 1.26* | 1.42* | 1.18* |
| Starting 607/REL607 | 1.10 | 1.25* | 1.27* |
| Starting 606/Starting 607 | 1.26* | 1.09* | 0.94 |

\*Heterogeneous two sample two-tail t-test significant, $P < 0.05$ .

**Figure S 2:**
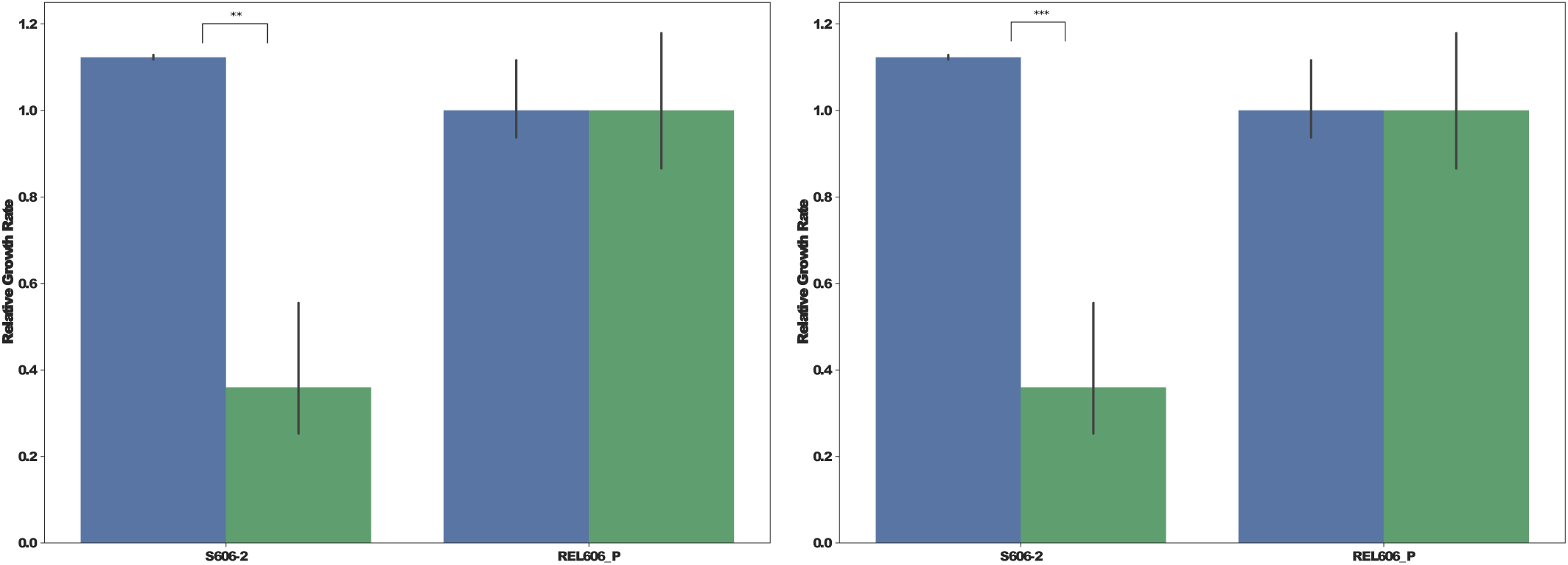
Dependence of the relative growth rate on the environment previously experienced. The left side shows our final REL606-derived clones and the right side shows our final REL607-derived clones. The blue bars correspond to their growth at 15°C after being passed in 43°C. The green bars correspond to their growth at 15°C after having grown at 15°C. In both cases, the strains have been normalized to their starting strain’s average growth rate. ** Significant at *α* = 0.01, *** significant at *α* = 0.001 for a heterocedastic two samples one-tailed t-test.

**Table S 5:** Top loadings of the PCs for the ancestral strains.

| PC1 | PC2 | PC3 | PC4 |
| --- | --- | --- | --- |
| Sodium Butyrate | Nalidixic Acid | pH5 | Niaproof 4 |
| D-Saccharic Acid | Potassium Tellurite | Aztreonam | Sodium Butyrate |
| L-Glutamic Acid | Fusidic Acid | 8NaCl | D-Mannitol |
| L-Aspartic Acid | p-HydroxyPhenylacetic Acid | Keto-Glutaric Acid | D-Serine |
| Vancomycin | Minocycline | D-Serine | D-Mannose |
| N-Acetyl--DMannosamine | Sodium Bromate | Propionic Acid | pH5 |
| L-Malic Acid | Propionic Acid | 4NaCl | trehalose |
| Fusidic Acid | Aztreonam | Niaproof 4 | 1NaCl |
| Glucuronamide | D-Raffinose | 1NaLactate | D-Fructose |
| Keto-Glutaric Acid | Lincomycin | Sodium Butyrate | D-Galactose |
| D-Fructose6-PO4 | D-Fructose6-PO4 | Troleandomycin | D-Melibiose |
| Troleandomycin | Keto-Glutaric Acid | Lithium Chloride | D-Saccharic Acid |
| L-Lactic Acid | Acetic Acid | Sodium Bromate | a-D-Lactose |
| b-Methyl-DGlucoside | L-Alanine | D-Saccharic Acid | Vancomycin |
| L-Alanine | Glucuronamide | 1NaCl | maltose |

**Figure S 3:**
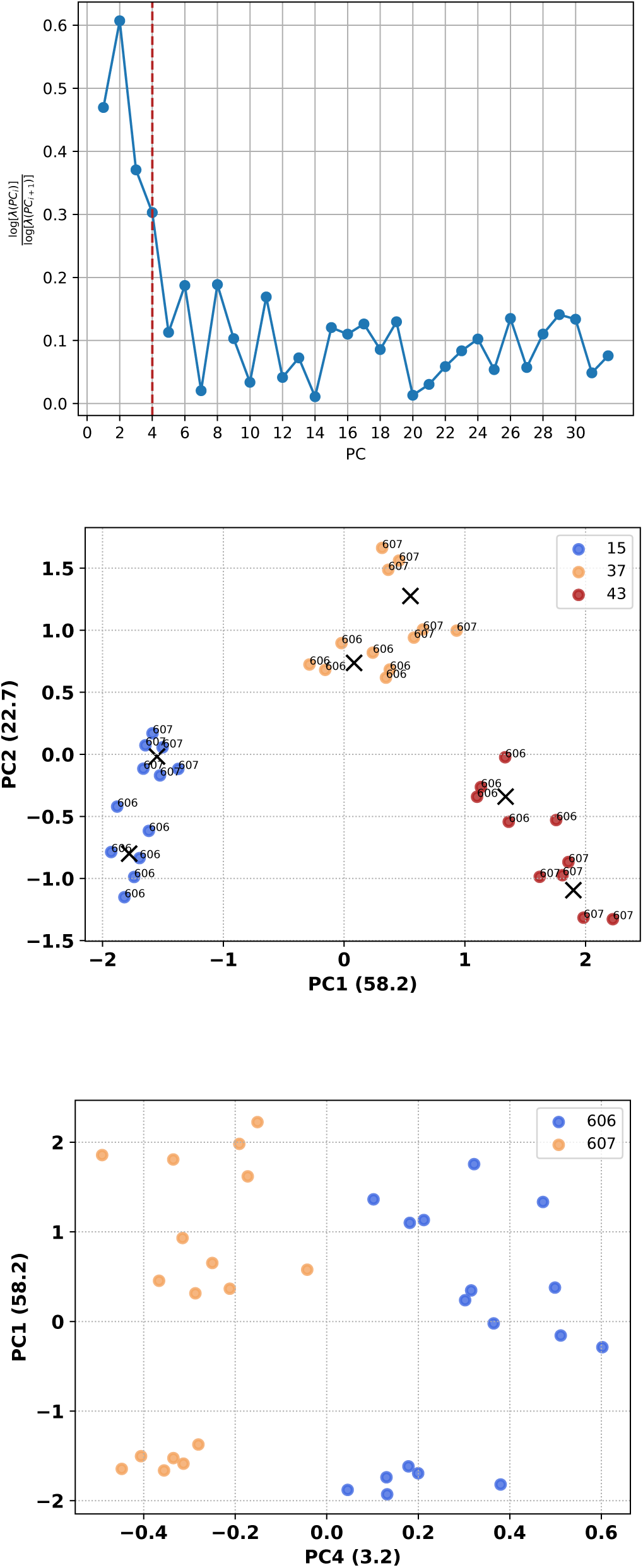
Principal component analysis of the phenotypic fingerprint of our starting strains of our laboratory evolution experiment at three temperatures. Top, the talus plot which shows random oscillations beyond 4 principal components, indicating that these are sufficient to explain the structure of the data. The red line marks the number of components that have signal in the data. Middle, the data projected on the first and second principal components. Bottom, the fourth principal component carries information about the strain.

**Table S 6:** Top loadings of the PCA of the 606 linage

| PC1 | PC2 | PC3 | PC4 |
| --- | --- | --- | --- |
| Glucuronamide | p-HydroxyPhenylacetic Acid | Nalidixic Acid | Pectin |
| D-Saccharic Acid | Keto-Glutaric Acid | Lithium Chloride | Amino-Butyric Acid |
| N-Acetyl--DMannosamine | Propionic Acid | Guanidine HCl | "Hydroxy-D,L-Butyric Acid |
| Keto-Glutaric Acid | 4NaCl | Fusidic Acid | Niaproof 4 |
| L-Aspartic Acid | D-Serine | Vancomycin | L-Arginine |
| L-Glutamic Acid | Niaproof 4 | Sodium Butyrate | D-Lactic Acid Methyl Ester |
| L-Malic Acid | Keto-Butyric Acid | Propionic Acid | L-Galactonic Acid Lactone |
| Vancomycin | D-Saccharic Acid | Niaproof 4 | Keto-Butyric Acid |
| D-Fructose6-PO4 | pH5 | Minocycline | Methyl Pyruvate |
| L-Galactonic Acid Lactone | D-Fructose6-PO4 | Aztreonam | L-Aspartic Acid |
| Sodium Butyrate | Sodium Butyrate | Keto-Butyric Acid | Potassium Tellurite |
| Fusidic Acid | L-Alanine | Potassium Tellurite | D-Gluconic Acid |
| L-Alanine | Guanidine HCl | D-Serine | Citric Acid |
| Aztreonam | D-Glucose6-PO4 | Sodium Bromate | Minocycline |

**Table S 7:**
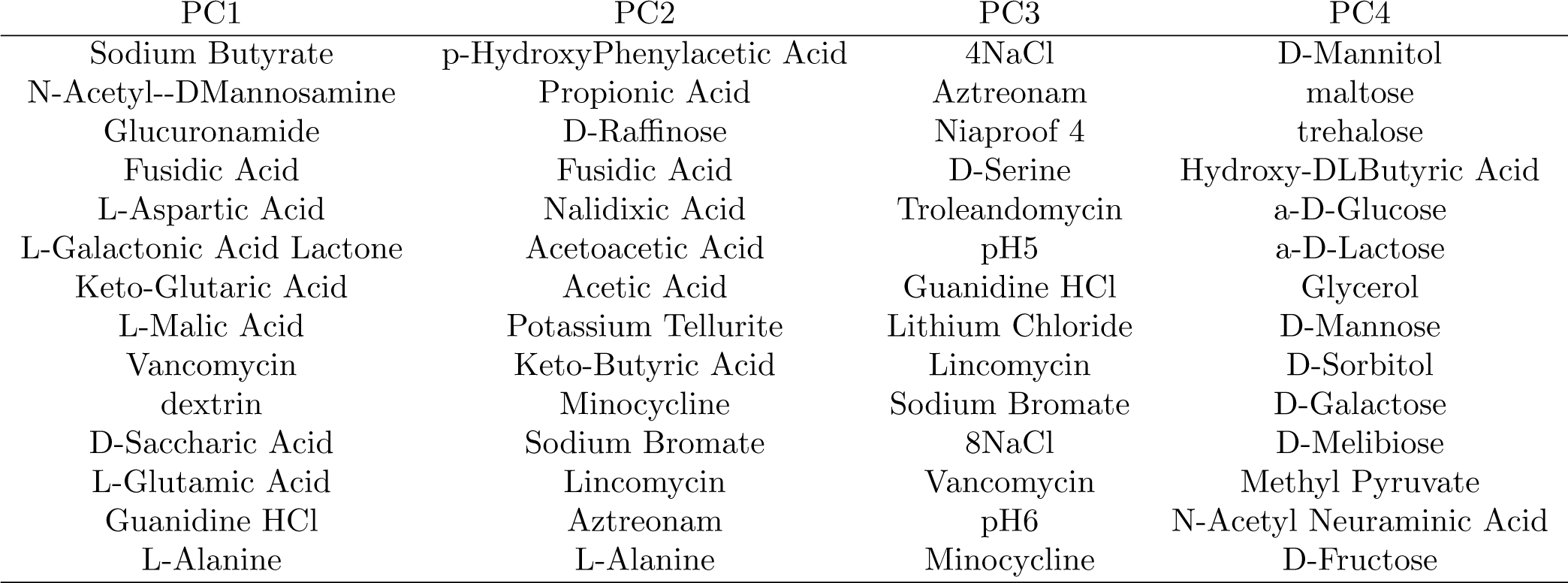
Top loadings of the PCA of the 607 linage

| PC1 | PC2 | PC3 | PC4 |
| --- | --- | --- | --- |
| Sodium Butyrate | p-HydroxyPhenylacetic Acid | 4NaCl | D-Mannitol |
| N-Acetyl--DMannosamine | Propionic Acid | Aztreonam | maltose |
| Glucuronamide | D-Raffinose | Niaproof 4 | trehalose |
| Fusidic Acid | Fusidic Acid | D-Serine | Hydroxy-DLButyric Acid |
| L-Aspartic Acid | Nalidixic Acid | Troleandomycin | a-D-Glucose |
| L-Galactonic Acid Lactone | Acetoacetic Acid | pH5 | a-D-Lactose |
| Keto-Glutaric Acid | Acetic Acid | Guanidine HCl | Glycerol |
| L-Malic Acid | Potassium Tellurite | Lithium Chloride | D-Mannose |
| Vancomycin | Keto-Butyric Acid | Lincomycin | D-Sorbitol |
| dextrin | Minocycline | Sodium Bromate | D-Galactose |
| D-Saccharic Acid | Sodium Bromate | 8NaCl | D-Melibiose |
| L-Glutamic Acid | Lincomycin | Vancomycin | Methyl Pyruvate |
| Guanidine HCl | Aztreonam | pH6 | N-Acetyl Neuraminic Acid |
| L-Alanine | L-Alanine | Minocycline | D-Fructose |

**Figure S 4:**
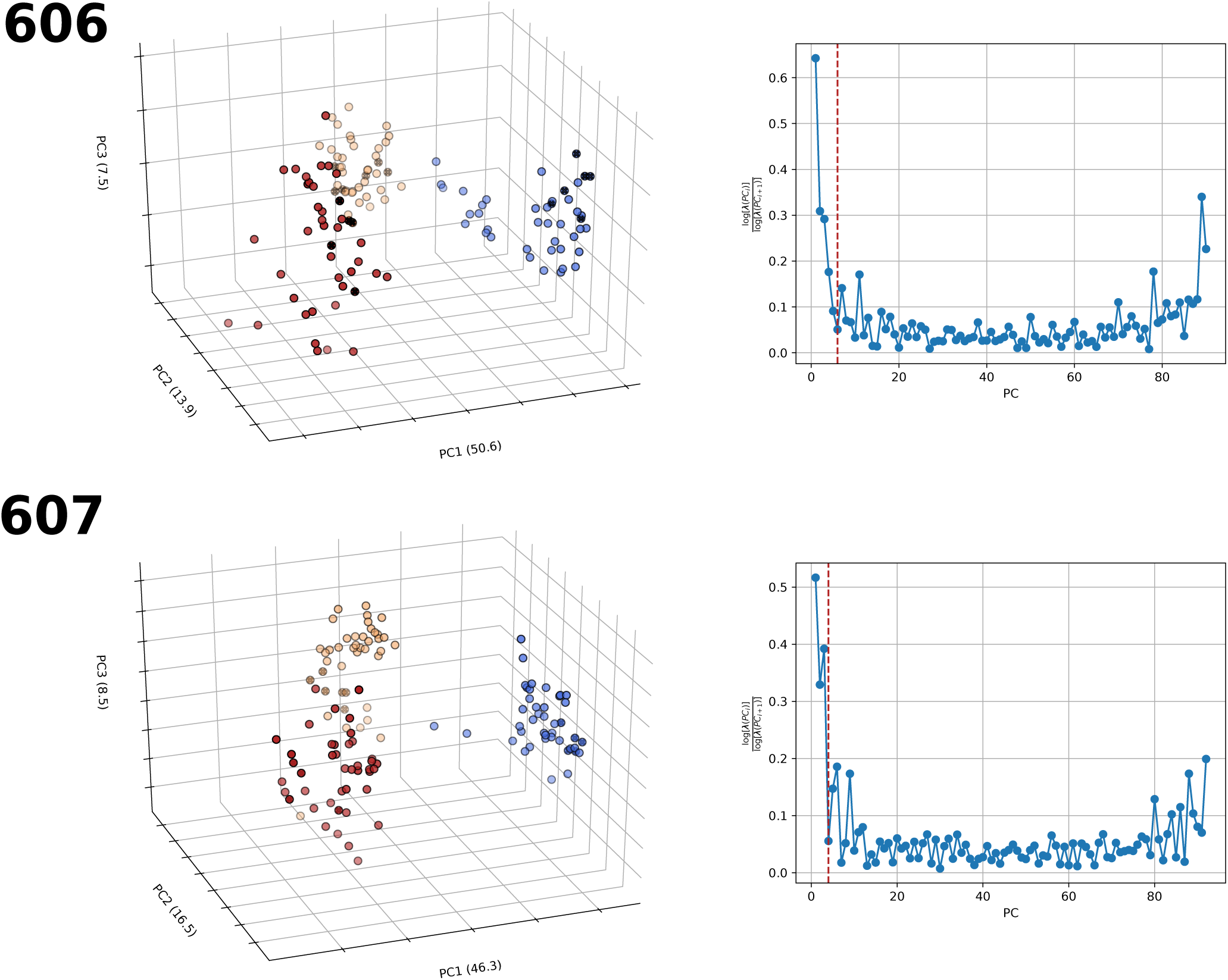
First three principal components and talus plot. The sample points are colored by temperature. In the talus plot, the red line marks the number of components that have signal in the data.

**Figure S 5:**
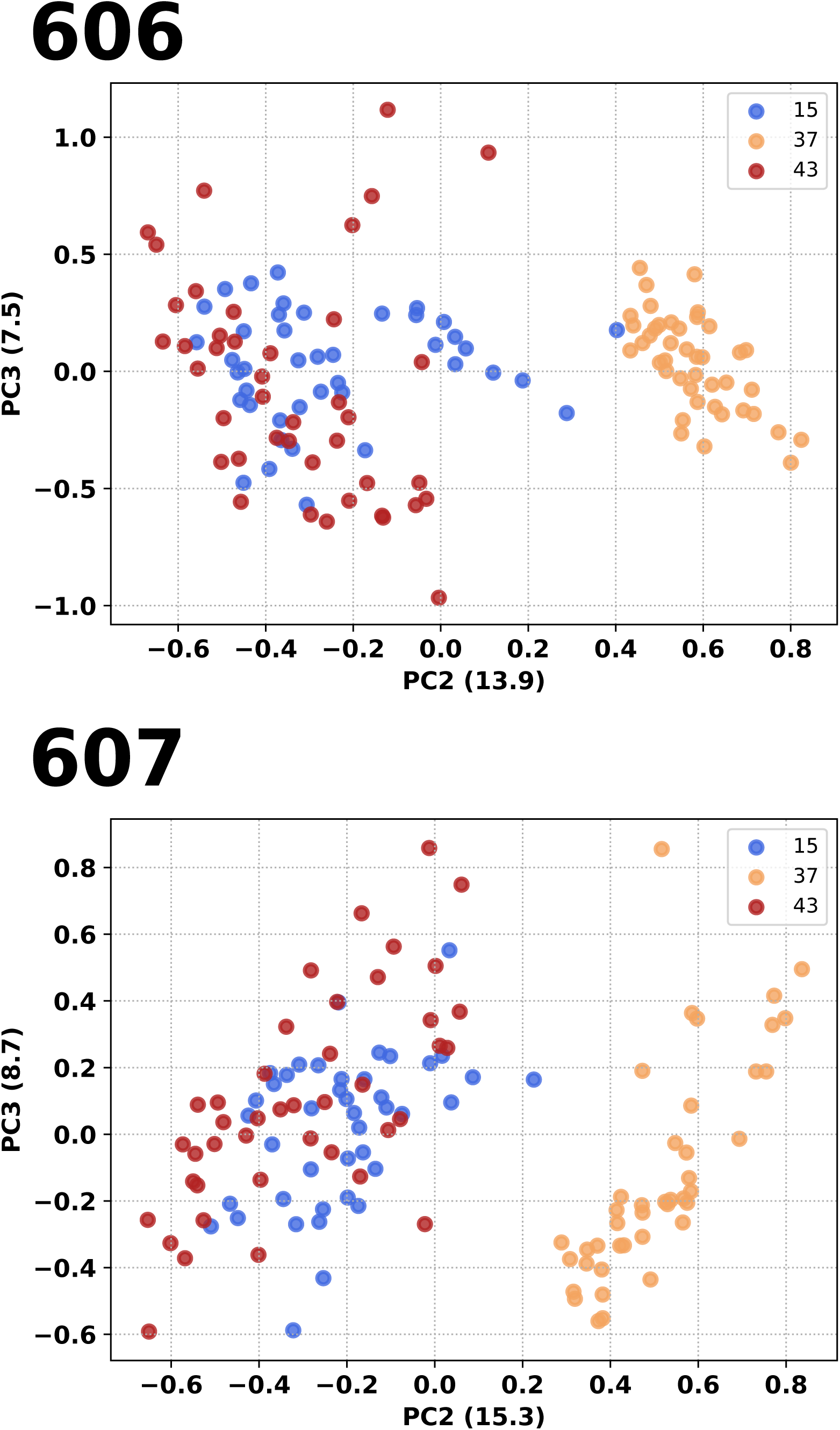
The second principal component separates the perturbed temperatures from the optimal. The sample points are colored by temperature.

**Table S 8:** Summary of the direction of our final clones’ evolved phenotypic response with respect to their starting strain.

| Strain | PC1 |  | PC2 |  | PC3 |  | PC4 |  |
| --- | --- | --- | --- | --- | --- | --- | --- | --- |
| | $T = 15$ | $T = 43$ | $T = 15$ | $T = 43$ | $T = 15$ | $T = 43$ | $T = 15$ | $T = 43$ |
| F606-1 | Unrestored | Unrestored | Unrestored | Unrestored | Uninformative | Uninformative | Partially Restored | Uninformative |
| F606-2 | Partially Restored | Reinforced | Unrestored | Reinforced | Uninformative | Over-restored | Restored | Uninformative |
| F606-3 | Unrestored | Unrestored | Unrestored | Unrestored | Restored | Over-restored | Partially Restored | Uninformative |
| F606-4 | Unrestored | Unrestored | Unrestored | Partially Restored | Uninformative | Unrestored | Partially Restored | Novel |
| F607-1 | Unrestored | Partially Restored | Partially Restored | Unrestored | Uninformative | Uninformative | Unrestored | Reinforced |
| F607-2 | Partially Restored | Partially Restored | Reinforced | Partially Restored | Novel | Uninformative | Reinforced | Reinforced |
| F607-3 | Partially Restored | Partially Restored | Partially Restored | Unrestored | Uninformative | Uninformative | Unrestored | Reinforced |
| F607-4 | Unrestored | Partially Restored | Unrestored | Partially Restored | Novel | Uninformative | Unrestored | Reinforced |
| R606-1 | Partially Restored | Unrestored | Unrestored | Unrestored | Uninformative | Uninformative | Restored | Uninformative |
| R606-2 | Partially Restored | Partially Restored | Partially Restored | Partially Restored | Restored | Unrestored | Restored | Novel |
| R606-3 | Partially Restored | Unrestored | Restored | Unrestored | Uninformative | Unrestored | Uninformative | Uninformative |
| R606-4 | Unrestored | Reinforced | Unrestored | Partially Restored | Restored | Uninformative | Restored | Uninformative |
| R607-1 | Partially Restored | Partially Restored | Partially Restored | Partially Restored | Uninformative | Uninformative | Reinforced | Reinforced |
| R607-2 | Unrestored | Partially Restored | Partially Restored | Unrestored | Uninformative | Uninformative | Unrestored | Unrestored |
| R607-3 | Unrestored | Partially Restored | Unrestored | Unrestored | Uninformative | Uninformative | Reinforced | Reinforced |
| R607-4 | Unrestored | Unrestored | Unrestored | Unrestored | Uninformative | Uninformative | Reinforced | Unrestored |
| S606-1 | Partially Restored | Unrestored | Unrestored | Unrestored | Restored | Uninformative | Uninformative | Uninformative |
| S606-2 | Unrestored | Unrestored | Reinforced | Partially Restored | Restored | Unrestored | Partially Restored | Novel |
| S606-3 | Partially Restored | Partially Restored | Unrestored | Unrestored | Restored | Restored | Restored | Uninformative |
| S606-4 | Partially Restored | Reinforced | Partially Restored | Unrestored | Restored | Over-restored | Restored | Novel |
| S607-1 | Partially Restored | Unrestored | Partially Restored | Partially Restored | Uninformative | Novel | Partially Restored | Unrestored |
| S607-2 | Unrestored | Uninformative | Partially Restored | Partially Restored | Uninformative | Uninformative | Unrestored | Reinforced |
| S607-3 | Unrestored | Partially Restored | Unrestored | Partially Restored | Uninformative | Novel | Reinforced | Reinforced |
| S607-4 | Partially Restored | Partially Restored | Partially Restored | Partially Restored | Novel | Novel | Unrestored | Reinforced |

## Notes

Support was provided by NIH MSTP training grant T32-GM007288 to DB, the Fulbright program to XP, and NIH R01-CA164468-01 and R01-DA033788 to AB. The authors thank Cameron Smith, Daniel Martinez, Bill Jacobs, Brian Wreinwrick, Adel Malek, Yves Robert Juste, Andrea Madrid, Raymond Puzio and Max Gottesman for their contribution to the development of this project.

